# Convergence of Wnt, Growth Factor and Trimeric G protein signals on Daple

**DOI:** 10.1101/137125

**Authors:** Nicolas Aznar, Ying Dunkel, Nina Sun, Kendall Satterfield, Fang He, Inmaculada Lopez-Sanchez, Majid Ghassemian, Debashis Sahoo, Irina Kufareva, Pradipta Ghosh

**Author notes:** ***Current address***: Center of Infectious Diseases, West China Hospital and Division of Infectious Diseases, State Key Laboratory of Biotherapy, Sichuan University, Chengdu, Sichuan, PR China. **To whom correspondence should be addressed**: N.A and P.G, **Nicolas Aznar, Ph.D**; Department of Medicine, University of California, San Diego School of Medicine, La Jolla, California 92093-0651., **Pradipta Ghosh, M.D**; Professor, Departments of Medicine and Cellular and Molecular Medicine, University of California, San Diego School of Medicine; George E. Palade Laboratories for Cellular and Molecular Medicine, 9500 Gilman Drive, Room 333; La Jolla, California 92093-0651.

## Abstract

Cellular proliferation, differentiation, and morphogenesis are shaped by multiple signaling cascades; their concurrent dysregulation plays an integral role in cancer progression and is a common feature of many malignancies. Three such cascades that contribute to the oncogenic potential are the Wnt/Frizzled(FZD), growth factor-receptor tyrosine kinases (RTKs), and G-proteins/GPCRs. Here we identify Daple, a modulator of trimeric G-proteins and a Dishevelled (Dvl)-binding protein as an unexpected point of convergence for all three cascades. Daple-dependent activation of Gαi and enhancement of non-canonical Wnt signals is not just triggered by Wnt5a/FZD to suppress tumorigenesis, but also hijacked by growth factor-RTKs to stoke tumor progression. Phosphorylation of Daple by both RTKs and non-RTKs triggers Gαi activation and potentiates non-canonical Wnt signals that trigger epithelial-mesenchymal transition. In patients with colorectal cancers, concurrent upregulation of Daple and the prototype RTK, EGFR, carried poor prognosis. Thus, this work defines a novel growth factor↔G-protein↔Wnt crosstalk paradigm in cancer biology.

## Introduction

Molecular characterization of tumors has revealed that multiple signaling pathways are often simultaneously dysregulated in cancer cells. Although each of these pathways are often conceptualized as independent entities, their complex crosstalk shapes many aspects of cancers, e.g., proliferation, invasion, immune evasion, chemoresistance and stemness (Anastas, 2015; Bertrand et al, 2012; Morris & Huang, 2016; Peng et al, 2015; Rota & Wood, 2015; Song et al, 2015; Yanai et al, 2008). Aberrant Wnt/Frizzled, G proteins/G protein-coupled receptors [GPCRs] and growth factor/ receptor tyrosine kinase [RTK]-based signaling cascades are three examples of such pathways that are frequently dysregulated in cancers, and the crosstalk between these pathways are of paramount importance in driving several properties of cancer cells. For example, aberrant activation of β-catenin–dependent Wnt signals [the so-called canonical β-catenin–TCF/LEF transcriptional program] secondary to adenomatous polyposis coli (APC), axin, and β-catenin gainof-function mutations are associated with the development of colon cancer, desmoid tumors, gastric cancer, hepatocellular carcinoma, medulloblastoma, melanoma, ovarian cancer, pancreatic cancer, and prostate cancer [reviewed in (Morris & Huang, 2016)]. However, these mutations alone do not account for the observed β-catenin hyperactivity; instead, it is the crosstalk between the growth factor RTK and the β-catenin–dependent Wnt/Frizzled pathways that synergistically hyperactivate the β-catenin-dependent transcriptional program (Anastas, 2015; Nelson & Nusse, 2004). This growth factor/RTK↔Wnt/β-catenin crosstalk is a well-defined paradigm that is frequently encountered in cancers and allows growth factor RTKs to potentiate β-catenin signaling via distinct mechanisms [reviewed in (Anastas, 2015; Nelson & Nusse, 2004)]: **a)** by triggering PI3K-Akt signals, which in turn can inhibit the downstream kinase, GSK3β; inhibition of GSK3β prevents proteasomal degradation of β-catenin and results in increased accumulation of β-catenin, followed by nuclear localization; activated Akt can also directly phosphorylate β-catenin and enhance its transcriptional activity (Fang et al, 2007); **b)** by triggering the MAPK/ERK kinase cascade, which can stabilize β-catenin by evading proteasomal degradation (Ding et al, 2005; Kim & Choi, 2007; Lemieux et al, 2015; Yun et al, 2005); and finally, **c)** by increased shedding of β-catenin from Ecadherin-bound junctional complexes (Hazan & Norton, 1998; Lu et al, 2003). These mechanisms underscore the importance of concurrent aberrant signaling [triggered by sequential genetic/epigenetic ‘hits’]; in solid tumors, aberrations in as few as three driver genes/pathways appear to suffice for a cell to evolve into an advanced cancer (Vogelstein & Kinzler, 2015).

Although the elaborate crosstalk between growth factors and the Wnt/β-catenin-dependent signaling pathway is well-documented, nothing is known about how growth factors affect non-β-catenin–dependent Wnt signaling. The non-canonical Wnt pathway behaves as a double-edged sword; it suppresses tumorigenesis in normal epithelium and in early tumors, but also serves as a critical driver of epithelial-mesenchymal transition [EMT] and cancer invasion (Ara et al, 2016; Asem et al, 2016; Avasarala et al, 2013; Chen et al, 2011; Ford et al, 2014; Katoh, 2011; Katoh & Katoh, 2007; Qi et al, 2014; Tseng et al, 2016; Vela et al, 2014; Yan et al, 2016). We recently defined a novel paradigm in Wnt signaling in which Frizzled receptors (FZDs) activate the G proteins and trigger non-canonical Wnt signaling via Daple (CCDC88C), which is a Dishevelled (Dvl)-binding protein (Aznar et al, 2015). Daple directly binds Wnt5a-activated FZDs, and serves as a guanine nucleotide exchange factor (GEF) that activates the G protein, Gαi (Figure 1A). Upon ligand stimulation, Daple-GEF dissociates from Dvl, binds and displaces Dvl from FZDs (Figure 1B), assembles Daple-Gαi complexes (Figure 1C), and triggers the activation of trimeric Gi near ligand-activated FZDs. Activation of Gαi by Daple-GEF suppresses cAMP, whereas released ‘free’ Gβγ heterodimers enhance Rac1 and PI3K-Akt signals (Figure 1A). We and others have shown that Daple-dependent enhancement of non-canonical Wnt signals can suppress tumor growth (Aznar et al, 2015), but can also fuel EMT, trigger cancer cell migration and invasion (Aznar et al, 2015; Ishida-Takagishi et al, 2012) and drive metastasis (Ara et al, 2016). Furthermore, elevated expression of Daple-GEF in circulating tumor cells prognosticates a poor outcome (Barbazan et al, 2016). In doing so, Daple behaves like a double-edged sword-- a tumor suppressor early during oncogenesis, but mimics an oncogene and fuels metastatic invasion later. Who/what triggers this switch, was unclear.

**Figure 1:**
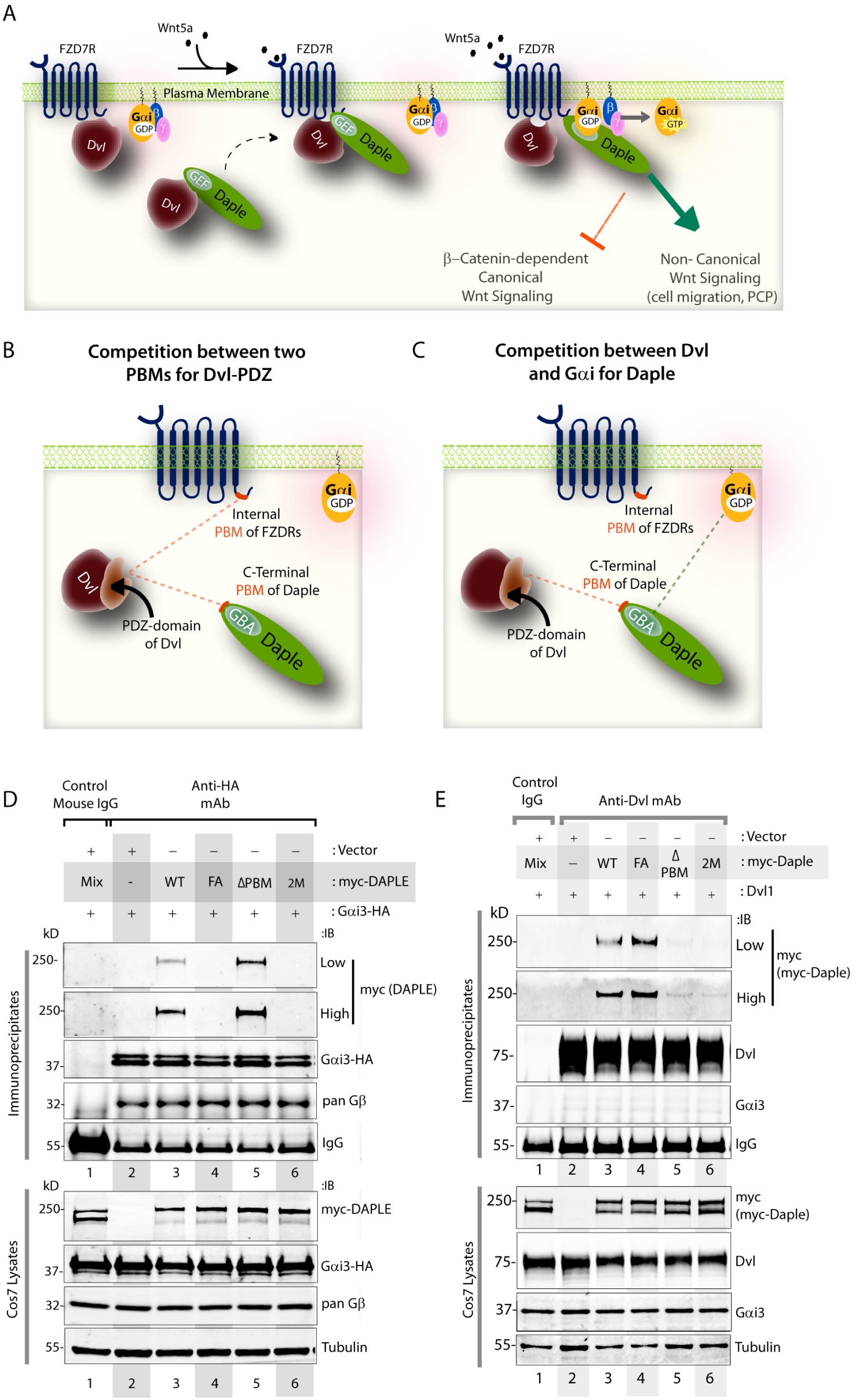
The <u>G</u>αi-<u>b</u>inding and <u>a</u>ctivating [GBA] and <u>P</u>DZ-<u>b</u>inding <u>m</u>otifs [PBM] in Daple allosterically inhibit each other’s functions. **(A)** Schematic summarizing how Daple enhances non-canonical Wnt signaling downstream of FZDs. (From left to right) In the absence of Wnt5a ligand, inactive FZD7Rs remain at the PM in complex with a PM-localized pool of Dvl, whereas Daple, via its PDZ-binding motif, remains in the cytosol in complex with cytosolic Dvl; G proteins at the PM exist as inactive Giαβγ trimers. Upon ligand [Wnt5a] stimulation, Dvl-Daple complexes dissociate, Daple is recruited to the cytoplasmic tails of activated receptors where it displaces Dvl and favors the assembly of receptor-Gαi complexes via its GBA motif. Using the same GBA motif Daple triggers the activation of Gαi within these complexes to antagonize β-catenin– dependent and trigger non-canonical Wnt signaling pathways. **(B-C)** Schematic showing two key intermolecular interplays encountered during non-canonical Wnt signaling via Daple, as shown in (Aznar et al, 2015). The first interplay is between FZD7R PBM and Daple PBM for binding to the PDZ-domain of Dvl (**B**). The second interplay is between Dvl and Gαi for binding to two distinct modules on Daple; Dvl binds to the PBM of Daple, whereas Gαi binds to the GBA motif on Daple (**C**). **(D)** Binding of full-length Daple to Gαi3 is increased upon deletion of Daple’s PBM motif (ΔPBM). Gαi3 was immunoprecipitated from equal aliquots of lysates of Cos7 cells coexpressing Gαi3-HA and myc-Daple [wild-type; WT, F1675A (GBA-deficient, FA), ΔPBM, or double mutant (FA+ΔPBM; a.k.a, 2M)] using anti-HA mAb and protein G beads. Immune complexes were analyzed for Daple (myc) and Gαi3 (HA) by immunoblotting (IB). Low and high exposures are shown for Daple. Gβ was monitored as a positive control for Gαi3-bound proteins. Quantification of blots is shown in Figure 1-figure supplement 1A. **(E)** Binding of full-length Daple to Dvl is increased upon disabling Daple’s GBA motif. Dvl2 was immunoprecipitated from equal aliquots of lysates of Cos7 cells coexpressing Dvl and myc-Daple [WT and mutants as in **D**] with anti-Dvl2 mAb and subsequently with protein G beads. Immune complexes were analyzed for Daple (myc) and Dvl by immunoblotting (IB). Low and high exposures are shown for Daple. Quantification of blots is shown in Figure 1-figure supplement 1B.

Here we show that growth factor- RTKs could trigger such switch by hijacking the Daple-Gαi cascade. Phosphorylation of Daple by both RTKs and non-RTKs enhances Daple’s ability to bind and activate Gαi and potentiate non-canonical Wnt signals that trigger epithelial-mesenchymal transition. Thus, Daple serves as an unexpected point of convergence for three major signaling cascades, Wnt/FZD, G proteins, and RTKs, whose concurrent dysregulation plays an integral role in cancer progression. Findings also reveal how aberrant growth factor signaling may reprogram a tumor suppressive pathway to potentiate non-canonical Wnt signals that drive EMT during cancer progression.

## Results and Discussion

### Allosteric regulation of Daple’s G protein regulatory function by its C-terminal PDZ-binding motif

We recently showed (Aznar et al, 2015) that upon Wnt5a stimulation, cytosolic Daple:Dvl complexes dissociate, and instead Daple:Gαi3 complexes are assembled. In vitro protein-protein interaction assays using purified recombinant proteins had revealed that binding of the PDZ domain of Dvl to a C-terminal PDZ-binding motif [PBM] in Daple could compete with the binding of Gαi to the <u>G</u>αi-<u>b</u>inding and <u>a</u>ctivating [GBA] module of Daple (Figure 1C); increasing concentration of Gαi displaced Daple from Dvl. Such competition was unexpected because the GBA [aa 1665-1685] and the PBM [aa 2025-2028] modules in Daple are separated by ~350 aa and are within a disordered stretch of the molecule with no semblance to known 3D-structural modules, suggesting that the observed competition between Dvl:Daple-PBM and Gαi:Daple-GBA interactions may be allosteric. When we analyzed Dvl:Daple and Gαi:Daple interactions in cells using co-immunoprecipiation assays, we found that binding of Daple to Gαi was consistently higher (increased ~2 fold) when the PBM module of Daple was deleted [Daple-ΔPBM, a mutant that cannot bind Dvl (Oshita et al, 2003)] (Figure 1D; Figure 1-figure supplement 1A). Conversely, binding of Daple to Dvl was consistently higher (increased ~2 fold) when the GBA module of Daple was disabled by a single point mutation [Daple-F1675A, a mutant that cannot bind Gαi; (Aznar et al, 2015)] (Figure 1E; Figure 1-figure supplement 1B). These findings suggest that the PBM and GBA modules of Daple allosterically inhibit each other. Because the increase in binding of Daple-ΔPBM to Gαi3 was also seen in *in vitro* pulldown assays in which the G protein was bacterially expressed (Figure 1-figure supplement 1C-D), we conclude that allosteric inhibition of Daple:Gαi interaction by Daple’s PBM is likely to be due to change in the properties of Daple, and not Gαi. Although the structural basis for such inhibition remains unknown, based on these findings and our prior results (Aznar et al, 2015) we conclude that the Dvl-PDZ:Daple-PBM and Gαi:Daple-GBA interactions antagonistically inhibit each other. Binding of Dvl to Daple may therefore suppress Daple•Gαi-dependent enhancement of FZD/non-canonical Wnt signaling. These findings agree with others’ observations that overexpression of Dvl can indeed suppresses FZD/G protein-dependent non-canonical Wnt signaling (Kilander et al, 2014). Release of such suppression, perhaps by specific events, may trigger the initiation/enhancement of Daple-dependent G protein signaling in cells.

### Multiple tyrosine kinases phosphorylate Daple’s PBM

We hypothesized that post-translational modifications at the Dvl-PDZ:Daple-PBM interface may disrupt the interface, and thereby serve as a trigger for a shift in Daple’s preferred binding partner from Dvl to Gαi, and for the initiation of Daple:Gαi-dependent non-canonical Wnt signaling in cells. Daple’s C-terminal PBM [<u>Y</u>^2023^E<u>Y</u>^2025^GCV^2028^-COOH] is notable for the presence of two evolutionarily conserved tyrosines (Figure 2A). Multiple kinase prediction programs (Scansite Motif Scan, MIT; NetPhos 2.0, Denmark; KinasePhos, Taiwan; Phospho-Motif Finder, HPRD) indicate that RTKs and non-RTKs may phosphorylate Daple at those tyrosines. We carried out in vitro kinase assays using wild-type [WT] and non-phosphorylatable [in which Tyr, Y2025, either alone (YF), or along with Y2023 (Y2F), is mutated to Phe, F] mutants of His-tagged purified Daple-CT [aa 1650-2028] protein using various recombinant TKs. We found that while Y2025 can be phosphorylated by all RTKs tested (EGFR, PDGFR, InsR) as well as the non-RTK Src, Y2023 is phosphorylated exclusively by Src (Figure 2B-C). MALDI MS/MS spectra of phosphorylated peptides [aa 1650-2028; this segment contains a total of 5 tyrosines] confirmed that while RTKs like EGFR phosphorylated Daple at Y1750 and Y2025, Src phosphorylated Daple at tyrosines Y1655, Y2023 and Y2025; only one of them, Y1699 has been identified as a phosphosite in cells using high-throughput phosphoproteomics (HTP) <u>(www.Phosphosite.org)</u> (Figure 2-figure supplement 1). Using WT and a non-phosphorylatable mutant of Daple (in which Y2025 is mutated to Phe, F [YF]) and a generic anti-pan-pTyr (pY) antibody for immunoblotting, we confirmed that Y2025 is indeed phosphorylated in cells responding to EGF (Figure 2D), as well as in cells expressing constitutively active Src (SrcCA; Figure 2E). Using a combination of Daple-WT, Daple-ΔPBM [which lacks Y2025], Daple-YF, and another non-phosphorylatable mutant in which both Y2023 and Y2025 were mutated to F [Y2F] we further confirmed that Src indeed phosphorylated Daple at both tyrosines, Y2025 and Y2023 (Figure 2C; Figure 2- figure supplement 2). Finally, we generated an antibody against pY2023 and pY2025 and confirmed that it can detect Daple exclusively upon phosphorylation *in vitro* and in cells (Figure 2- figure supplement 3). Using this antibody, we further confirmed that endogenous Daple is phosphorylated at Y2023 and Y2025 in cells responding to EGF (Figure 2F). We conclude that multiple RTKs and the non-RTK Src phosphorylate Daple at Y2025 within the PBM, and that Src can also phosphorylate Daple at Y2023.

**Figure 2:**
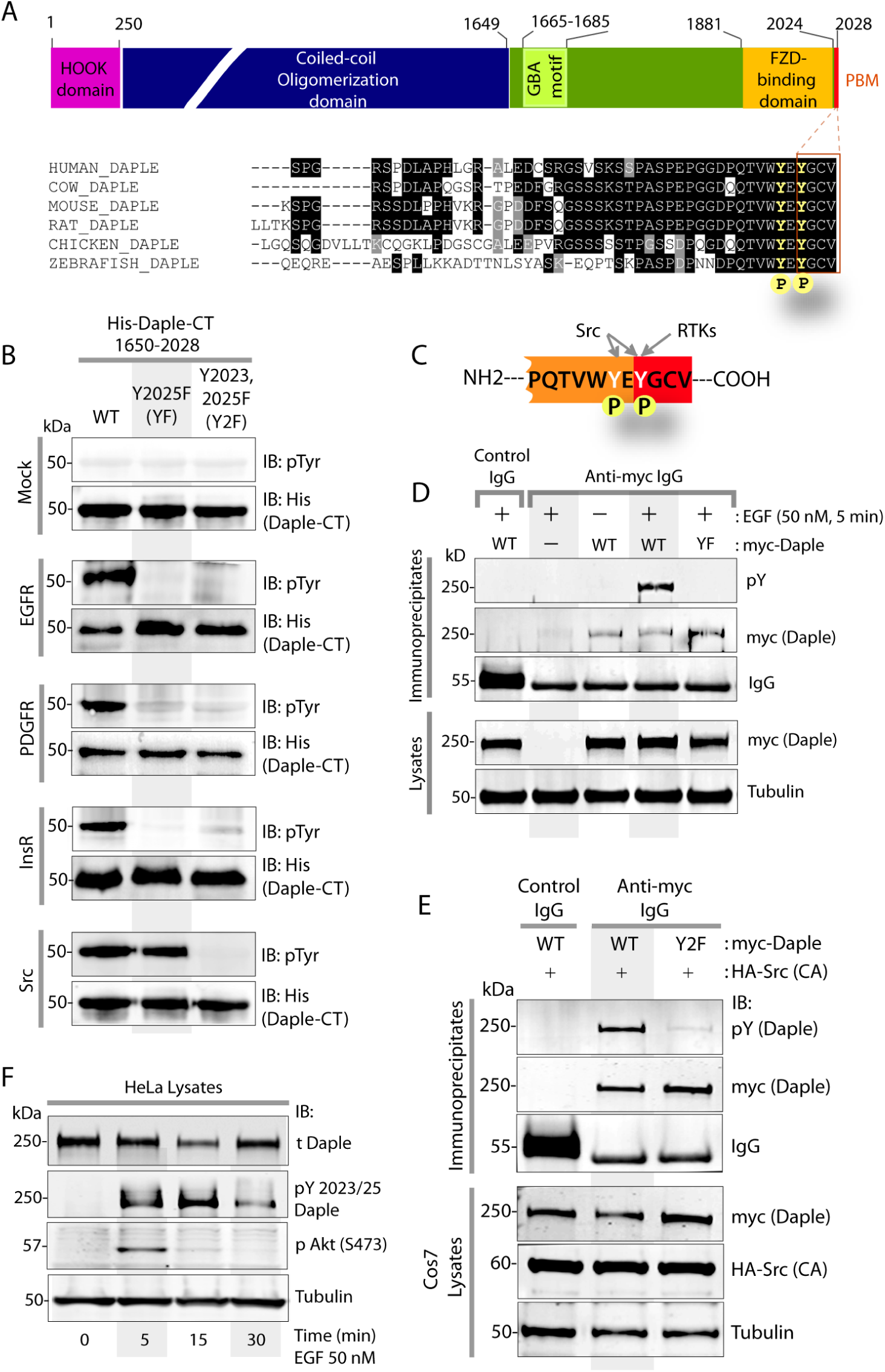
Multiple TKs phosphorylate Daple’s C-terminal PBM module. (**A**) *<u>Top</u>:* Schematic showing the domain arrangement of Daple. *<u>Bottom</u>*: An alignment of Daple’s C-terminus in various species showing an evolutionary conserved PDZ-binding motif [PBM; red box]. Two tyrosines, one within the PBM and one just proximal to the PBM are highlighted in yellow. (**B-C**) Both RTKs and the non-RTK, Src phosphorylate Daple’s PBM. In vitro kinase assays were carried out using several recombinant kinases (~ 50 ng) and equal aliquots (~3 μg) of wild-type (WT) or non-phosphorylatable mutants [Y2025F (YF) and Y2023/2025F (Y2F)] His-Daple CT proteins as substrate. Entire reactions were separated by SDS PAGE and analyzed by immunoblotting with anti-pTyr mAb. RTKs like EGFR, PDGFR and InsR primarily phosphorylated Daple-CT-WT, but phosphorylation was virtually abolished for both YF and Y2F mutants. Src phosphorylated Daple-CT-WT, and to a lesser extent the YF mutant; phosphorylation was virtually abolished in Y2F mutant. Schematic representation of the findings shown in **B**. (**D**) EGF triggers tyrosine phosphorylation of Daple’s PBM in cells. Daple was immunoprecipitated from equal aliquots of lysates of Cos7 cells co-expressing EGFR and myc-Daple [WT or YF mutant] using anti-myc or control IgG. Bound complexes were analyzed for phosphorylated and total Daple using a pan-pTyr (pY) and myc antibodies, respectively. Phosphorylation was detected in Daple-WT, but not the non-phosphorylatable YF mutant. (**E**) Src phosphorylates Daple’s PBM in cells. Daple was immunoprecipitated from equal aliquots of lysates of Cos7 cells co-expressing a constitutively active (CA; Y527F) Src mutant and myc-Daple [WT or Y2F mutant] using anti-myc or control IgG. Bound complexes were analyzed for tyrosine phosphorylated Daple as in **D**. Phosphorylation was detected in Daple-WT, but not the non-phosphorylatable Y2F mutant. (**F**) EGF triggers tyrosine phosphorylation of endogenous Daple in cells. HeLa cells were serum starved overnight [0.2% FBS] prior to stimulation with 50 nM EGF for the indicated time prior to lysis. Equal aliquots of lysates were analyzed by immunoblotting (IB) for tyrosine phosphorylated Daple using a phospho-specific anti-Daple (pY2023, pY2025) antibody [see Methods]. Phosphorylation of Daple coincides with the onset of Akt phosphorylation and peaks at 15 min after EGF stimulation.

### Daple is required for activation of Gαi and enhancement of Akt and Rac1 signals downstream of EGFR

Next we asked how the newly discovered phosphoevent on Daple may impact EGF signaling. We hypothesized that if tyrosine phosphorylation triggers a switch of Daple-bound complexes from Dvl to G protein, some of the G protein regulatory functions of Daple we reported previously, e.g., activation of Gαi and enhancement of Akt and Rac1 signals (Aznar et al, 2015) must be impacted in cells responding to growth factors. To answer that, first we asked if Daple and its ability to bind and activate Gαi3 was essential for cellular response to EGF, the ligand for the prototype RTK, EGFR. We found that Daple is essential for EGF response in a manner similar to what we reported previously in the case of Wnt5A/FZD7 (Aznar et al, 2015)— First, using an anti-Gαi•GTP mAb that specifically recognizes Gαi in a GTP-bound active conformation (Lane et al, 2008) and previously validated by us to study Daple-GEF (Aznar et al, 2015), we found that compared to controls Daple-depleted cells failed to enhance Gαi3 activity in response to EGF (Figure 3A; Figure 3-figure supplement 1), indicating that Daple is required for such activation. Second, Daple-depleted cells also failed to enhance Rac1 activity (Figure 3B-C) and Akt phosphorylation (Figure 3D-E).

**Figure 3:**
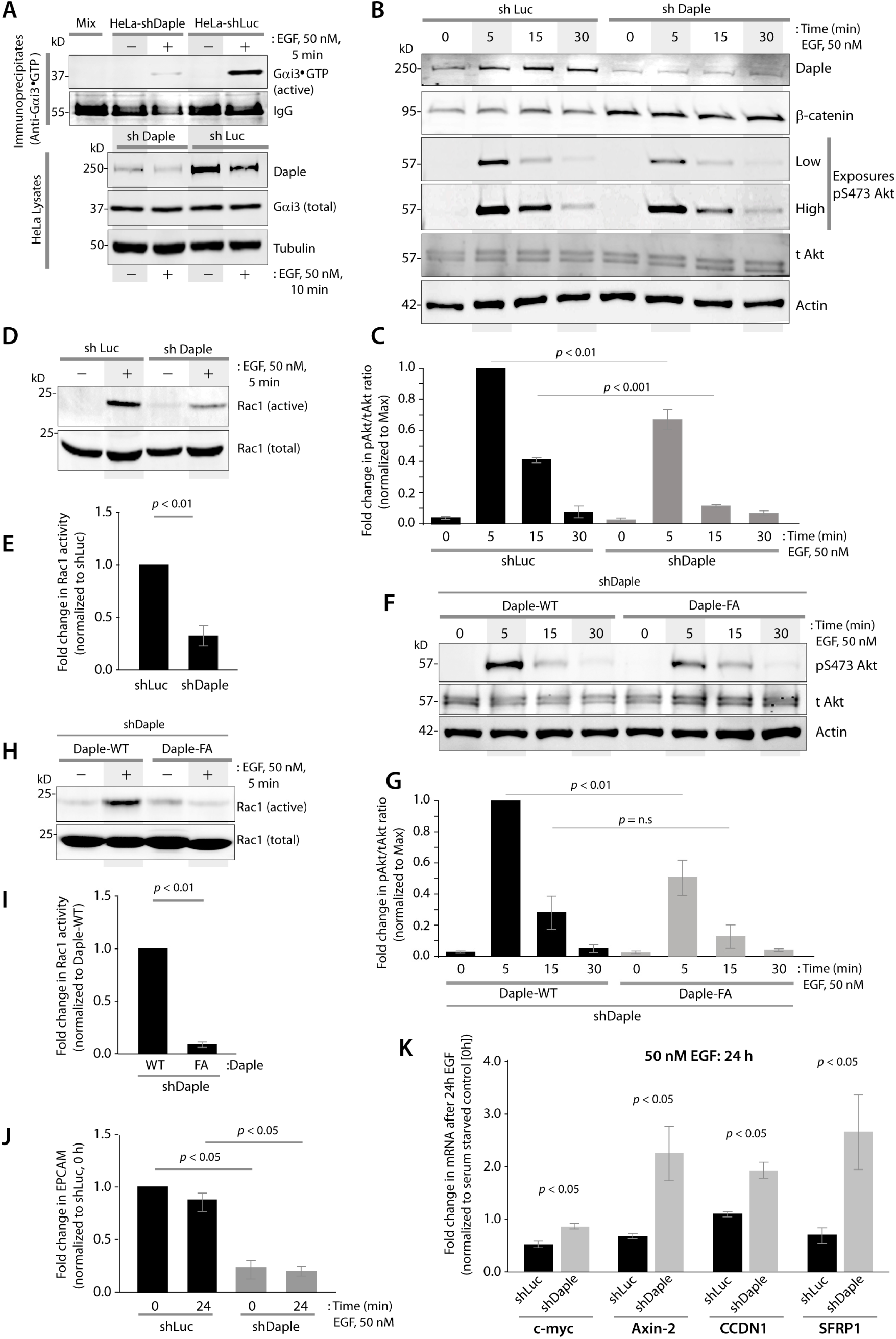
Daple is required for activation of Gαi3, Rac1 and Akt signals and in the antagonistic inhibition of β-catenin–dependent Wnt signals downstream of EGF/EGFR. **(A)** Control (shLuc) and Daple-depleted (shDaple) HeLa cells were serum-starved (0.2% FBS) and treated (+) or not (−) with EGF for 10 min prior to lysis. Equal aliquots of lysates were subjected to immunoprecipitation with conformational mAbs that selectively recognize active GTP-bound Gαi-subunits. A 1:1 mix [MIX] of each lysate was incubated with control IgG (negative control). Immune complexes (top) and lysates (bottom) were analyzed for active Gαi3•GTP and total Gαi3 by immunoblotting (IB). EGF stimulation robustly activates Gαi3 in control cells; the extent of activation is reduced in Daple-depleted cells. Quantification of blots is shown in Figure 3-figure supplement 1. (**B, C**) Compared to control (shLuc) cells, activation of Rac1 in response to EGF is impaired in Daple-depleted (shDaple) HeLa cells. Analysis for Rac1 activation by pulldown assays using GSTPBD (see *methods*). Active GTP-bound Rac1 was assessed by immunoblotting (**B**). Bar graphs (**C**) display the quantification of Rac1 activation. Error bars represent mean ± S.D of three independent experiments. (**D, E**) Compared to control (shLuc) cells, phosphorylation of Akt induced by EGF is reduced in Daple-depleted (shDaple) HeLa cells. Serum-starved (0.2% FBS) HeLa cells stimulated with EGF were analyzed for Akt phosphorylation at S473 by immunoblotting (**D**). Bar graphs (**E**) display quantification of phosphorylated/total Akt. Error bars represent mean ± S.D of three independent experiments. (**F, G**) The GBA motif of Daple is required for the enhancement of Akt phosphorylation in cells responding to EGF. Activation of Rac1 was assessed by GST pulldown and immunoblotting as in **B**. Compared to Daple-WT expressing cells, activation of Rac1 by EGF is impaired in cells expressing Daple-F1675A (FA) mutant (**F**). Bar graphs (**G**) display quantification of Rac1 activation. Error bars represent mean ± S.D of three independent experiments. (**H, I**) Akt phosphorylation at S473 in response to EGF was assessed in HeLa cells expressing Daple-WT or DapleFA as in **D**. Compared to those expressing Daple-WT, phosphorylation of Akt is reduced in cells expressing Daple-F1675A (FA) mutant (**H**). Bar graphs (**I**) display quantification of phosphorylated/total Akt. Error bars represent mean ± S.D of three independent experiments. (**J**) Daple is required for the suppression of canonical Wnt target genes induced by EGF. Control (shLuc) and Daple-depleted (shDaple) HeLa cells were analyzed for C-MYC, AXIN-2, CCDN1 and SFRP-1 mRNA by qPCR. Bar graphs display the fold change in each RNA (y axis) after EGF stimulation normalized to their respective condition without stimulation. Error bars represent mean ± S.D of three independent experiments.

Next we interrogated if the ability to activate Gαi was necessary for Daple to enhance the Akt and Rac1 pathways downstream of EGFR. For this we used previously characterized (Aznar et al, 2015) HeLa cells that are depleted of endogenous Daple and stably express either WT or a GEF-deficient Daple-F1675A (FA) mutant at levels close to endogenous levels. Upon EGF stimulation, HeLa-Daple-WT, but not Daple-FA cells enhanced Rac1 (Figure 3F-G) and Akt (Figure 3H-I). When it came to markers of EMT, we previously showed that compared to Daple-WT, Daple-FA cells have significantly lower levels at baseline of expression of Vimentin (VIM), Lysyl-oxidase (LOXL3) and the EMT-inducing transcriptional repressor ZEB1 (Aznar et al, 2015). We found that such holds true also for the adhesion molecule, EPCAM (Figure 3J), which can act as a double-edged sword during metastasis (van der Gun et al, 2010). Because none of these markers showed ligand(EGF)-dependent changes above the baseline changes, these findings suggest that reversible regulation of Daple’s PBM may not have long term effects on the transcription of the EMT target genes above and beyond the maximal steady-state effects already attributable to the presence or absence of Daple.

Finally, we analyzed the effect of Daple on EGF stimulated changes in canonical Wnt→β-catenin/TCF/LEF target genes. We also found that the expression of several β-catenin/TCF/LEF target genes were enhanced in Daple-depleted cells (Figure 3K), indicating that Daple antagonizes the β-catenin-dependent Wnt pathway in a manner similar to what we observed previously downstream of Wnt5A. Taken together, these studies indicate that much like the Wnt5A/FZD pathway (Aznar et al, 2015), Daple and its G protein modulatory function is required for several components of signaling also downstream of EGF/EGFR. We hypothesized, tyrosine phosphorylation of Daple’s PBM may serve as an acute trigger for the Daple→Gαi axis by simply disrupting the Daple-Dvl interaction and releasing the allosteric inhibition imposed by Dvl.

### Tyrosine phosphorylation abolishes Daple’s ability to bind Dvl

To investigate the consequences of tyrosine phosphorylation of Daple, we first asked if the binding specificity between Dvl-PDZ and Daple-PBM is controlled by the newly identified phosphoevents within/flanking Daple’s PBM. We found that binding of His-Daple-CT to GST-Dvl2-PDZ was reduced when the Daple-CT protein was phosphorylated *in vitro* with EGFR (Figure 4A). Similarly, when myc-Daple was phosphorylated in cells by co-expressing the catalytically active wild-type Src kinase, binding of Src-phosphorylated full length Daple to GST-Dvl-PDZ was also reduced (Figure 4B, D). By contrast, binding of Src-phosphorylated full-length Daple to GST-Gαi3 was increased (Figure 4C, D). To study the effect of phosphorylation of Daple-PBM in cells, we generated phosphomimicking and non-phosphorylatable mutants of full length myc-Daple in which the multi-RTK substrate site, Y2025 was replaced with either glutamate (YE) or phenylalanine (YF), respectively. Additional mutants were created in which both Y2025 and Y2023 were substituted (Y2E, or Y2F) to study the effect of Src-dependent phosphorylation on those residues. We found that both phosphomimicking myc-Daple mutants, YE and Y2E, failed to bind GST-Dvl-PDZ (Figure 4E; Figure 4- figure supplement 1A-B). Unexpectedly, we observed that the nonphosphorylatable YF mutant of Daple also bound poorly to Dvl (~60% less than WT; Figure 4E; Figure 4- figure supplement 1B) suggesting that the residue Y2025 may play an important role in the interface between Daple and Dvl that cannot be fulfilled by Phe [although both Y and F feature a 6-carbon aromatic ring, the latter lacks a hydroxyl(-OH) group which imparts Y its polar characteristics and allows participation in hydrogen(H)-bonds and improved protein stability (Pace et al, 2001)]. To gain structural insights into these findings, we generated a 3D homology-based model of Daple’s PBM bound to Dvl’s PDZ domain using previously solved structures of Dvl2-PDZ domain co-crystallized with various peptides from the Pocketome (Kufareva et al, 2012). The model showed that while both tyrosines are predicted to interact directly with Dvl by forming H-bonds [Y2025 with Asp (D)331 in Dvl, and Y2023 with Ser(S)286 in Dvl], the two tyrosines also stabilize each other with a third H-bond (Figure 4F). These findings explain why negative charge generated by phosphorylation at Y2025 (or Y2025E mutation) alone is sufficient to abolish Daple:Dvl interaction [likely to repel D331 on Dvl]. Because the interface is enriched in Tyr-OH mediated H-bonds, replacement of Y with F is indeed predicted to destabilize/weaken the Daple-PBM:Dvl-PDZ interface, consistent with findings in our binding assays (Figure 4A, B, E). These results are consistent with prior carefully conducted (Y→F) mutagenesis experiments which have documented the importance of Y-OH mediated interactions to the energy of protein-protein interactions (England et al, 1997).

**Figure 4:**
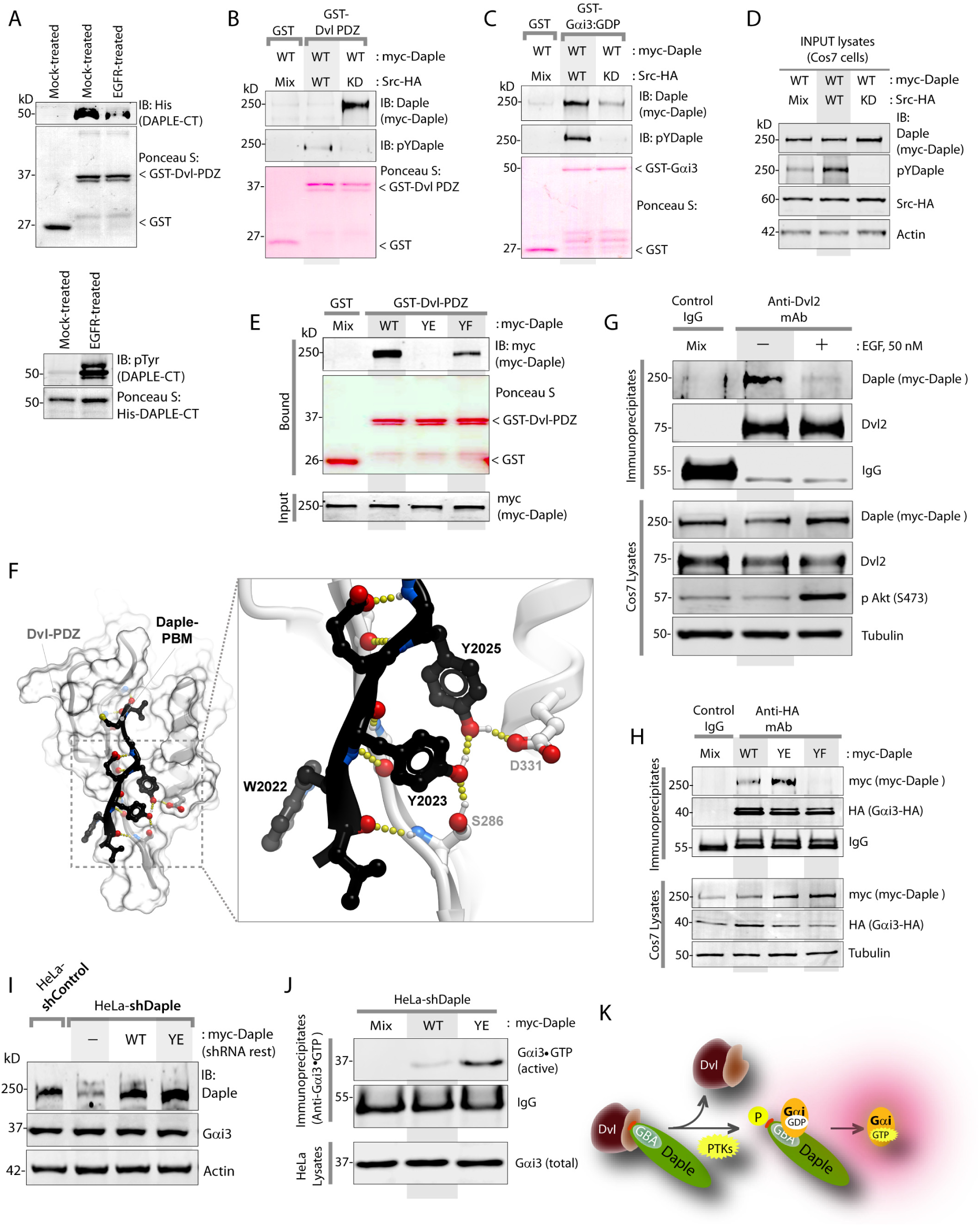
Tyrosine phosphorylation inhibits Daple’s ability to bind Dvl, but favors binding to and activation of Gαi. (**A**) Phosphorylation of Daple by EGFR inhibits binding to Dvl. His-Daple-CT proteins were either phosphorylated in vitro with recombinant EGFR kinase or mock-treated exactly as in Figure 2B prior to use in GST pulldown assays with GST-Dvl2-PDZ immobilized on glutathione Sepharose beads. *<u>Top</u>*: Bound proteins were analyzed for Daple (anti-His) by immunoblotting (IB). Equal loading of GST proteins was confirmed by ponceau S staining. *<u>Bottom</u>*: Equal loading of His-Daple-CT proteins and tyrosine phosphorylation by EGFR kinase was confirmed by ponceau S staining and immunoblotting with pTyr mAb. (**B-D**) Tyrosine phosphorylated Daple loses binding to Dvl and preferentially binds Gαi3. Full length Daple was phosphorylated in cells by Src kinase as in Figure 2E by coexpressing WT or kinase dead [KD] HA-Src with myc-Daple-WT. Equal aliquots of lysates was used as source of Daple in pulldown assays with GST-Dvl2-PDZ (**B**) or GST-Gαi3 (**C**) immobilized on glutathione Sepharose beads. Lysates (**D**) and bound proteins (**BC**) were analyzed for total (anti-Daple) and tyrosine phosphorylated Daple (pYDaple; pY2023/25 Daple) by immunoblotting (IB). (**E**) Lysates of Cos7 cells expressing full length myc-Daple [WT, phosphomimicking Y2025E (YE) or phosphodeficient Y2025F (YF) mutants] were used as sources of Daple in GST pulldown assays with GST-Dvl2-PDZ immobilized on glutathione Sepharose beads. *<u>Top</u>*: Bound proteins were analyzed for myc-Daple by immunoblotting with anti-myc IgG. *<u>Bottom</u>*: Equal expression of various myc-Daple constructs in lysates was confirmed by immunoblotting. Binding is virtually abolished in the case of Daple-YE and significantly reduced in the case of Daple-YF. Fold change in Daple binding to GST-Dvl2-PDZ is displayed as bar graph in Figure 4-figure supplement 1B. (**F**) Homology-based model of Daple’s PBM (black ribbon and sticks) bound to Dvl2-PDZ domain (grey surface) was generated using solved crystal structures of Dvl2-PDZ bound to multiple peptides [see *Methods*]. Both Y2023 and Y2025 in Daple’s PBM appear to partake in the stabilization of the Dvl2-Daple interface. Each tyrosine is predicted to form H-bonds (yellow dotted lines) with each other and with Dvl2 (D331 and S286). Replacing either Y with E is predicted to disrupt the interaction, whereas replacing either Y with F is predicted to significantly compromise the interaction by disrupting H-bonds. (**G**) EGF triggers the dissociation of Daple:Dvl complexes in cells. Dvl was immunoprecipitated from equal aliquots of lysates of serum starved Cos7 cells that were stimulated (+) or not (-) with 50 nM EGF prior to lysis. *<u>Top</u>*: Immune complexes were analyzed for Daple by immunoblotting. *<u>Bottom</u>*: Lysates were analyzed for equal expression of Dvl and Daple, and to confirm successful stimulation of Akt phosphorylation by EGF. Fold change in Daple binding to Gαi3 is displayed as bar graph in Figure 4-figure supplement 1C. (**H**) Gαi3 was immunoprecipitated from equal aliquots of lysates of Cos7 cells coexpressing myc-Daple [WT, phosphomimicking Y2025E (YE) or phosphodeficient Y2025F (YF) mutants] and Gαi3-HA using anti-HA IgG. *<u>Top</u>*: Immune complexes were analyzed for Daple (anti-myc) and Gαi3 (anti-HA) by immunoblotting. *<u>Bottom</u>*: Equal expression of various myc-Daple constructs and Gαi3-HA in lysates was confirmed by immunoblotting. (**I**) Generation of HeLa cell lines stably expressing various Daple constructs (see also *Methods).* Whole-cell lysates from control (shControl) or Daple-depleted (shDaple) HeLa cells stably expressing Daple-WT and YE were analyzed for Daple, Gαi3 and actin by immunoblotting (IB). Band densitometry (using LiCOR Odyssey) confirmed that various Daple constructs were expressed at levels ~1.0- 1.5 fold of endogenous. (**J**) Compared to Daple-WT, phosphomimicking Daple-Y2025E mutant (YE) enhances activation of Gai. Equal aliquots of lysates from Daple-depleted HeLa cells stably expressing Daple-WT or Daple-YE grown in 10% FBS were analyzed for Gαi-activity using conformational anti-Gαi3•GTP mAb exactly as in **3A**. These studies were done at steady-state because YE mutation represents constitutive phosphorylation at Y2025. Fold change in active:total Gαi3 is displayed as bar graph in Figure 4-figure supplement 1F. (**K**) Schematic summarizing the findings in panels **3A-J**. *<u>From left to right</u>*: Phosphorylation of Daple’s PBM by protein tyrosine kinases (PTKs) triggers dissociation of Daple:Dvl complexes and enables Daple to bind and activate Gαi.

To determine if these findings with *in vitro*-phosphorylated Daple or phosphomimicking YE mutants hold true in cells, we analyzed the abundance of Dvl:Daple complexes in cells before and after stimulation of the prototype RTK EGFR with EGF. Co-immunoprecipitation studies confirmed that the amount of Daple complexed with Dvl drops significantly (by ~75-80%) within 15 min after EGF stimulation (Figure 4G), like what was observed in response to Wnt5a ligand (Aznar et al, 2015). Taken together, these findings demonstrate that growth factors can indeed disrupt the Daple:Dvl interaction, and that tyrosine phosphorylation of Daple’s PBM by multiple RTKs and non-RTKs could serve as one such trigger.

### Tyrosine phosphorylation of Daple favors binding to and activation of Gαi downstream of EGF

Because binding of Dvl to Daple’s PBM allosterically inhibits Daple:Gαi interaction (Figure 1C) (Aznar et al, 2015), next we asked if the loss of Dvl:Daple interaction upon EGF stimulation enhances Daple’s ability to bind and activate Gαi. Consistent with our prior findings that tyrosine-phosphorylation of Daple favors interaction with G protein (Figure 4C), binding for YE was enhanced (~1.5 fold) and binding of the YF mutant was abolished (Figure 4H; Figure 4-figure supplement 1C). As for binding to FZD7, it appeared that tyrosine phosphorylation may have no effect because both Daple-WT and Daple-YE bound to a similar extent; however, Daple-YF bound less (Figure 4- figure supplement 1D-E). Finally, we confirmed that increased binding of Gαi to Daple-YE [observed in **4H**] indeed translated to increased G protein activity in cells because compared to cells expressing Daple-WT, activation of Gαi was higher in cells expressing the phosphomimicking Daple-YE mutant [which mimics constitutive phosphoactivation of Daple by PTKs; Figure 4I-J].

Thus, it appears that compared to Daple-WT, the phosphomimicking YE mutant did not bind Dvl, but bound and activated Gαi more efficiently, and had no effect at all on binding to FZD7 receptor. The non-phosphorylatable YF mutant, on the other hand, bound Dvl and FZD7, albeit with reduced affinity, but did not bind Gαi. These findings indicate that the two tyrosines in Daple’s PBM have a direct impact on binding to Dvl, and have allosteric impacts on two other key interactions mediated by distinct modules in Daple-CT that are located upstream of the PBM, i.e., FZD-binding domain and Gαi-binding and activating (GBA) motif (see Figure 2-figure supplement 1). Although the mechanisms of such allosteric effect remain unclear, phosphorylation of the two tyrosines in Daple-PBM downstream of growth factor RTKs abolishes Daple’s ability to bind Dvl, but increased its ability to bind and activate Gαi3 (Figure 4K).

### Tyrosine phosphorylation of Daple’s PBM enhances non-canonical Wnt signals, triggers EMT

We previously showed that Daple enhances non-canonical Wnt signals (Rac1 and PI3K/Akt signals), antagonistically suppresses β-catenin-dependent Wnt signals, inhibits proliferation/growth but triggers EMT and enhances tumor cell migration, all via activation of G proteins (Aznar et al, 2015). To study the impact of tyrosine-phosphorylation of Daple on these phenotypes we generated HeLa cell lines stably depleted of endogenous Daple and rescued by expressing Daple-WT or various YE/YF mutants at close to endogenous levels (Figure 5-figure supplement 1A). Because Y2025 is a common target of RTKs and non-RTKs and substitution of Y2025 with E is sufficient for maximally dissociating Daple:Dvl complexes and for binding and activating Gαi, we proceeded with in-depth characterization of the cells expressing Daple-WT and the YE mutant which mimics a constitutively phosphoactivated state and is sufficient for abolishing the Daple:Dvl interaction (Figure 4E) and augmenting Daple’s ability to bind and activate Gαi (Figure 4H-J). Because the non-phosphorylatable YF mutations continued to bind Dvl (Figure 4E) and FZD7 (Figure 4-figure supplement 1D-E), albeit weaker than Daple-WT, but abolished interaction with Gαi3 (Figure 4H), we cautiously analyzed this mutant as a non-phosphorylatable control that is likely to be nonresponsive to EGF stimulation.

Hyperactivation of Gαi in Daple-YE cells was associated with the enhancement of all previously published functions of Daple-GEF (Aznar et al, 2015). For example, compared to Daple-WT cells, Daple-YE cells displayed a greater antagonistic suppression of both levels of β-catenin protein (Figure 5A-B; Figure 5-figure supplement 1B-C) as well as levels of expression of its transcriptional targets (myc, CCND1 and SFRP1; Figure 5C**, left panel**). Surprisingly, two other targets of β-catenin, osteopontin [OPN; which is known as a master regulator of EMT (Kothari et al, 2016) via its ability to stabilize vimentin (Dong et al, 2016)] and Axin2 [AXIN2, which enhances EMT via induction of Snail (Yook et al, 2006)] were not suppressed, and instead were increased in Daple-YE cells (Figure 5C**, right panel**). Thus, constitutive phosphoactivation of Daple’s PBM mimicked in Daple-YE cells was associated with suppression of β-catenin/TCF/LEF transcriptional targets, but key targets that trigger EMT escaped such suppression. Suppression of β-catenin–dependent Wnt signaling in Daple-YE cells was accompanied also by reduced growth of these cells under both anchorage-dependent and -independent conditions (Figure 5D-E; Figure 5- figure supplement 1D-G). Compared to Daple-WT cells, Daple-YE cells displayed higher levels of Rac1 activity (Figure 5F) and Akt phosphorylation (Figure 5G), enhanced expression of markers of EMT (VIM and LOXL3; Figure 5H-I), and increased migration, as determined by transwell chemotaxis assays (Figure 5J). Daple-Y2E cells resembled the phenotypes observed in Daple-YE cells, only enhanced to a greater degree (Figure 5- figure supplement 1B-I). Cells expressing Daple-Y2F (a mutant that cannot bind Gαi, and binds poorly to Dvl and FZD) displayed an opposite phenotype; these cells showed increased colony growth and reduced migration (Figure 5- figure supplement 1B-J), and in doing so, resembled the previously characterized GEF-deficient Daple-F1675A mutant that cannot bind or activate Gαi (Aznar et al, 2015) (Figure 5- figure supplement 2). These findings are consistent with other instances in which imbalances in PDZ:PBM interactions perturb cellular homoeostasis and contribute to tumor cell phenotypes that ultimately fuel cancer progression [reviewed in (Subbaiah et al, 2011)].

**Figure 5:**
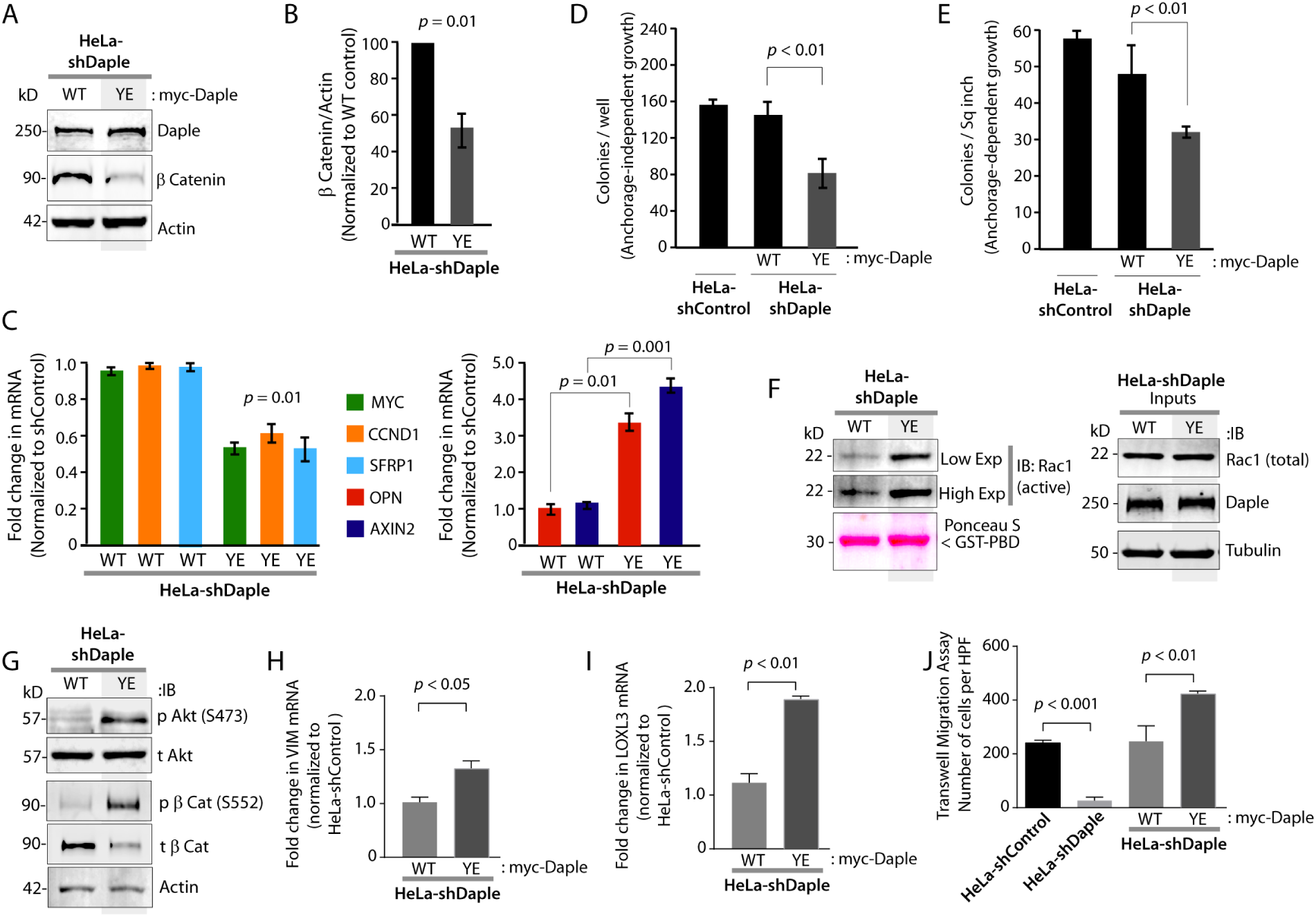
Tyrosine phosphorylated Daple enhances non-canonical Wnt signals, triggers EMT and cell migration. (**A-B**) Constitutive phosphoactivation of Daple [mimicked by Y2025E mutation] reduces the levels of β-catenin. Equal aliquots of lysates of Daple-depleted HeLa cells stably expressing Daple-WT or Daple-Y2025E (YE) were analyzed for β-catenin by immunoblotting (**A**). Daple and actin were monitored as loading controls. Bar graphs in **B** display the relative expression of β-catenin in **A**, normalized to actin and to Daple-WT cells. Error bars represent mean ± S.D of three independent experiments. (**C**) Constitutive phosphoactivation of Daple [mimicked by Y2025E mutation] supresses some, but not all canonical Wnt responsive β-catenin/TCF/LEF target genes. HeLa cell lines in **A** were analyzed by qPCR for the levels of mRNA for the indicated canonical Wnt target genes. Left and right panels display genes that are down- or up-regulated, respectively, in Daple-YE cells compared to Daple-WT. Results were normalized internally to mRNA levels of the housekeeping gene, GAPDH. Bar graphs display the fold change in each RNA (y axis) in cells expressing Daple-WT or Daple-YE normalized to the expression in shControl HeLa cells. Error bars represent mean ± S.D of three independent experiments. (**D-E**) Constitutive phosphoactivation of Daple [mimicked by Y2025E mutation] inhibits anchorage-independent (**D**) and anchorage-dependent (**E**) colony growth. Daple-WT and Daple-YE HeLa cell lines were analyzed for their ability to form colonies in soft agar (**D**) or on plastic plates (**E**) for 2–3 weeks (see also Figure 4-figure supplement 1D-G). For anchorage-independent growth in soft agar, the number of colonies was counted by light microscopy throughout the depth of the matrix in 15 randomly chosen fields. Bar graphs display the number of colonies (y axis) seen in each cell line. For anchorage-dependent growth, colonies were fixed and stained with crystal violet and counted using ImageJ (Colony counter). Bar graphs display the % inhibition of colony formation (y axis) seen in each condition normalized to shControl HeLa cells. (**F**) Constitutive phosphoactivation of Daple [mimicked by Y2025E mutation] enhances activation of Rac1. Daple-WT and Daple-YE HeLa cells grown in the presence of 10% FBS were analyzed for Rac1 activation by pulldown assays using GST-PBD. (**G**) Constitutive phosphoactivation of Daple [mimicked by Y2025E mutation] enhances Akt activation in HeLa cells, as determined by phosphorylation of Akt at S473, and increases phosphorylation of β-catenin at S552. Equal aliquots of lysates of Daple-WT and Daple-YE HeLa cells grown in the presence of 10% serum were analyzed for phospho (p) and total (t) Akt and β-catenin by immunoblotting (IB). (**H-I**) Constitutive phosphoactivation of Daple [mimicked by Y2025E mutation] triggers EMT. mRNA expression of the EMT markers, Vimentin and LOXL3, were analyzed in Daple-WT and Daple-YE HeLa cells by qPCR. Results were normalized internally to GAPDH. Bar graphs display the fold change in each mRNA (y axis) normalized to the expression in cells expressing vector control. Error bars represent mean ± S.D of three independent experiments. (**J**) Constitutive phosphoactivation of Daple [mimicked by Y2025E mutation] enhances cell migration. Control (shControl) and Daple-depleted (shDaple) cells and Daple-WT and Daple-YE HeLa cell lines were analyzed for their ability to migrate in transwell assays (see *Methods*). The number of migrating cells was averaged from 20 field-of view images per experiment. Representative images are shown in Figure 5-figure supplement 1H. Bar graphs display the number of cells/ high power field (HPF); data are presented as mean ± S.D; n = 3.

Selective enhancement of some EMT-associated β-catenin/TCF/LEF–dependent target genes (OPN and AXIN2) and suppression of other proliferation-associated transcriptional targets in Daple-YE cells (Figure 5C), despite the relatively lower levels of β-catenin in these cells (Figure 5A-B) was an unexpected observation. Such differential response of β-catenin/TCF/LEF target genes in Daple-YE cells responding to EGF could be due to differences in either the distribution or activity of the remaining pool of β-catenin. In fact, we found that enhanced Akt phosphorylation in Daple-YE cells (Figure 5G) is accompanied also by enhanced phosphorylation of β-catenin at Ser(S)552; phosphorylation by Akt at that site is known to enhance the transcriptional activity of β-catenin and promote tumor cell invasion (Fang et al, 2007). It is possible that higher activity of a smaller pool of β-catenin in Daple-YE cells is sufficient for targets like AXIN2 and OPN which are ultra-responsive to β-catenin/TCF/LEF (because of the large number of TCF sites in its promoter), but not sufficient for other targets. Additional factors (additional post-translational modifications, distribution, proteasomal degradation, protein-protein interactions) that affect the functions of β-catenin may coexist, and cannot be ruled out.

### Concurrent upregulation of Daple and EGFR in colorectal tumors carries poor prognosis

To determine the impact of crosstalk between growth factors and Daple-GEF on clinical outcome, we compared the mRNA expression levels to disease-free survival (DFS) in a data set of 466 patients with colorectal cancers (see *Methods*). Patients were stratified into negative (low) and positive (high) subgroups with regard to Daple (*CCDC88C)* and EGFR gene-expression levels with the use of the StepMiner algorithm, implemented within the Hegemon software (hierarchical exploration of gene-expression microarrays online; (Dalerba et al, 2011)] (Figure 6A). Kaplan-Meier analyses of DFS over time showed that among patients with high EGFR, expression of Daple at high levels carried a significantly poorer prognosis compared to those with low Daple (Figure 6B). Among patients with low EGFR, high expression of Daple was associated with a protective effect, although the trend was not statistically significant (Figure 6C). Conversely, among patients with high levels of Daple, survival was significantly different between those with high vs low EGFR (Figure 6D); no such trend was noted among patients with low Daple (Figure 6E). In fact, the high Daple/high EGFR signature carried a poorer prognosis compared to all other patients combined (Figure 6-figure supplement 1). Together, these results indicate that concurrent upregulation of the growth factor and non-canonical Wnt pathways spurred by Daple is associated with poorer clinical outcomes. Findings are in keeping with the fact that both pathways are frequently upregulated during oncogenic progression and are known to promote tumor cell dissemination (Aznar et al, 2015; Markman et al, 2010).

**Figure 6:**
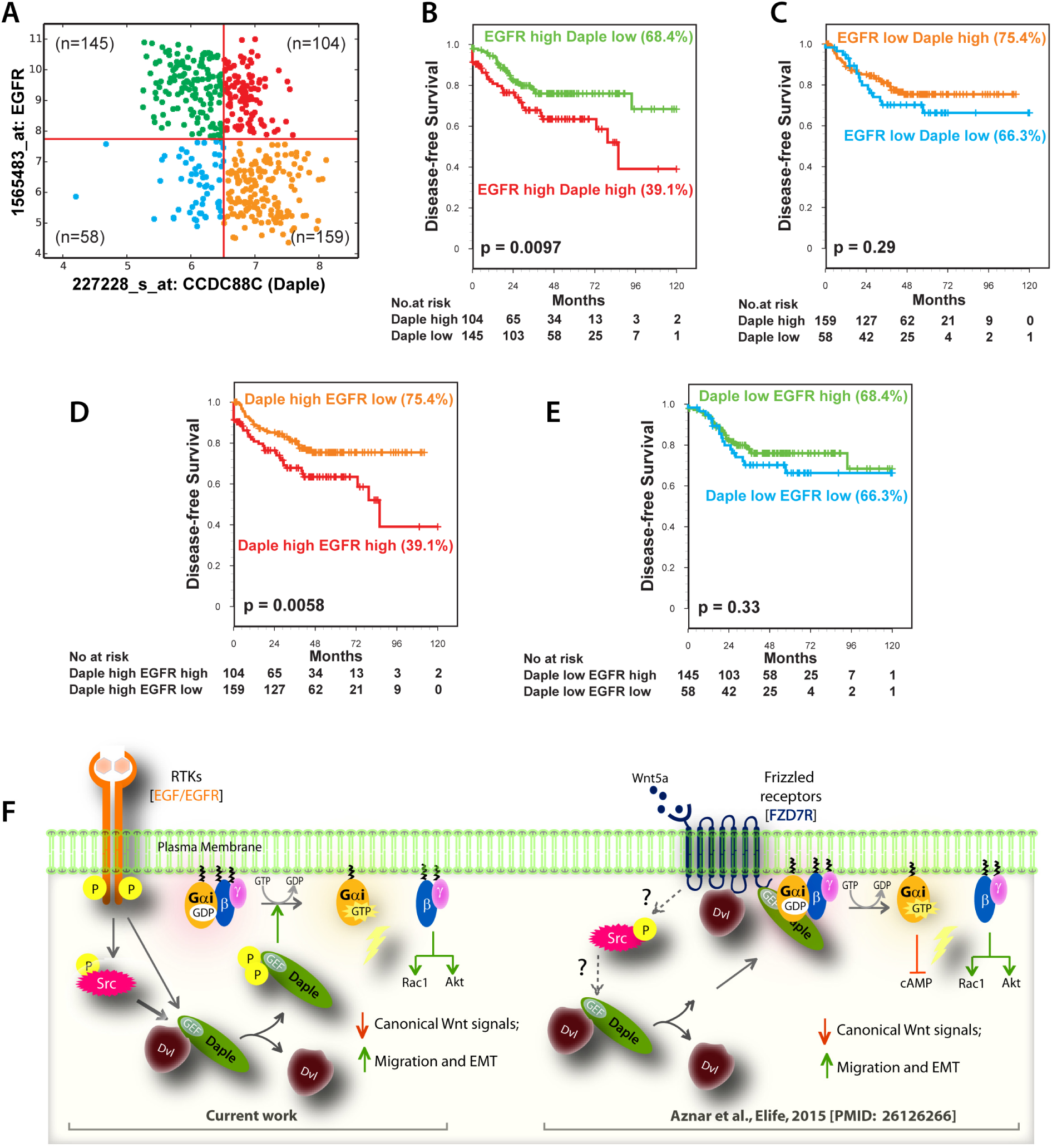
Concurrent expression of Daple (CCDC88C) and EGFR mRNA at high levels in colon cancer is associated with reduced disease-free survival (DFS). Hegemon software was used to graph individual arrays according to the expression levels of *EGFR* and *Daple(CCDC88C)* in a data set containing 466 patients with colon cancer (**A;** see *Methods*). (**B, C**) (**D, E**) Impact of varying levels of Daple expression within high *vs*. low EGFR groups. Survival analysis using Kaplan-Meier curves showed that among patients with high EGFR, concurrent expression of Daple at high levels carried significantly worse prognosis (**B**). Survival analysis among patients with low EGFR showed that high expression of Daple was somewhat protective, although it did not reach statistical significance (**C**). (**D, E**) Impact of varying levels of EGFR expression within high *vs*. low Daple groups. (**F**) Schematic of working model based on findings in current work [left] and prior work [right; (Aznar et al, 2015)]. *<u>Left</u>*: In the absence of growth factors or Src activation, Dvl remains complexed to Daple, and Gαi is largely inactive. Stimulation with growth factors like EGF triggers phosphorylation of Daple’s PBM [either directly by EGFR or indirectly via activation of the non-RTK Src]. Such phosphorylation triggers dissociation of Dvl:Daple complexes and favors the assembly of Daple:Gαi complexes and subsequent activation of Gαi. *<u>Right</u>*: Schematic summarizing the key events triggered by Wnt5a ligand demonstrated previously (Aznar et al, 2015). We previously showed that Daple-dependent activation of Gi leads to dissociation of the heterotrimer; Gαi and Gβγ subunits trigger signaling via their respective downstream intermediates (Rac1, PI3K, and cAMP). Both ligands, EGF (left) and Wnt5a (right) ultimately lead to antagonistic suppression of canonical Wnt signals and triggers EMT. Because Wnt5a is known to activate Src-family kinases (Shojima et al, 2015) it is possible that tyrosine-phosphorylation of Daple also plays a role in Wnt5a/FZD7R-dependent signaling via Daple.

## Conclusions

Previously we showed that non-canonical Wnt signals initiated upon binding of Wnt5a to FZDs are enhanced and propagated by the multimodular non-receptor GEF, Daple, via activation of Gαi proteins (Aznar et al, 2015). The major discovery in the current work is an alternate path for activation of Daple-dependent non-canonical Wnt signaling, one that is triggered by growth factors. In both instances, a key mechanistic step is that ligand stimulation [either Wnt5a (Aznar et al, 2015) or EGF (current work)] triggered the assembly of Daple:Gαi complexes to the detriment of Daple:Dvl complexes; such a switch in the composition of Daple-bound complexes was a pre-requisite for activation of G protein and enhancement of non-canonical Wnt signals through G protein intermediates [i.e., cAMP and ‘free’ Gβγ] (see legend, Figure 6F). Taken together, these results place Daple at the point of convergence between the Wnt/FZD, the growth factor RTKs, and G proteins/GPCRs, three major signaling mechanisms that are conventionally thought to operate independently. In doing so, this work defines a new paradigm in signal transduction, the first one of its kind at the cross-road of non-canonical Wnt and growth factor signaling pathways.

Mechanistically, the cross-talk between the three major pathways plays out at the level of Daple, Gαi and Dvl. Our experimental evidence indicates that Dvl and Gαi compete for binding Daple; such competition appears to be allosteric. By triggering tyrosine phosphorylation of Daple’s PBM, growth factors facilitate the dissociation of Daple:Dvl complexes and favor the assembly of Daple:Gαi complexes. These findings do not contradict the well-accepted notion that Dvl is required for non-canonical Wnt signaling, or that the Dvl-Daple interaction is required for Daple-dependent non-canonical Wnt signaling (Ishida-Takagishi et al, 2012). Because our findings specifically showed that the Dvl-Daple and Daple-G protein interactions antagonize each other, but both are essential for Daple-dependent non-canonical Wnt signaling cascade, we conclude that the reversible dynamic phosphoregulation of the Dvl-Daple interaction may be critical for proper signaling within this non-canonical cascade.

Our findings are also in keeping with the fact that many PDZ:PBM interactions between a variety of signaling molecules are known to be regulated by phosphoevents [reviewed in (Subbaiah et al, 2011)]. For example, multiple S/T kinases [like Bcr (Radziwill et al, 2003), CAMKII (Gardoni et al, 2003), PKA (Cohen et al, 1996)] are known to target either PDZs or PBMs. Although tyrosine phosphorylation of PBM has been reported in at least one instance, i.e., Syndecan-1 (SDC1), such phosphorylation did not impact binding to its PDZ partner, Tiam1 (Liu et al, 2013) [a key distinction from Daple is the position of Y in SDC1-PBM at next to the last aa from the end]. We conclude that disruption of Daple:Dvl binding by phosphoevents triggered by TKs is the first example of tyr-kinase-based signaling regulating a PDZ-PBM interaction and impacting G protein signaling. These insights pinpoint a clear ‘event’ that involves players from all three signaling pathways, i.e., TKs, Dvl and G proteins. Findings also add to the crosstalk between heterotrimeric G proteins and Dvl that have been reported previously-- Gβγ subunits released from Gα subunits can bind Dvl (Angers et al, 2006; Egger-Adam & Katanaev, 2010; Seitz et al, 2014), and target it for degradation (Jung et al, 2009). Further studies are warranted to assess the ramifications of this crosstalk.

Our findings using phosphomimicking (YE) and non-phosphorylatable (YF) tyrosine mutants of Daple in cell-based assays (Figure 5A-J) and hierarchical exploration of gene-expression microarrays on patient-derived colorectal tumors (Figure 6A-E) shed light onto how concurrent deregulation in growth factor and non-canonical Wnt signaling pathways may impact cancer progression. Although the major physiologic function of non-canonical Wnt signaling is in the establishment of planar cell polarity, tissue morphogenesis and suppression of tumors, non-canonical Wnt signaling is also known to promote the invasiveness and malignant progression of cancers (Sugimura & Li, 2010); the latter traits are also fuelled by growth factors (Lowery & Yu, 2012). Overexpression of Wnt5a has been found to be associated with aggressive tumor biology and poor prognosis (Huang et al, 2005; Kurayoshi et al, 2006; Pukrop & Binder, 2008). Deregulated growth factor signaling (e.g., copy number variations or activating mutations in RTKs, increased growth factor production/concentration) is also often encountered in advanced tumors (Lowery & Yu, 2012). We previously showed (Aznar et al, 2015) that Daple enhances non-canonical Wnt signaling via its ability to activate G-proteins downstream of Wnt5/ FZD7 receptors; it serves as a tumor suppressor in the normal epithelium and in early tumors (Aznar et al, 2015), but aids metastatic progression in advanced tumors and in circulating tumor cells (Ara et al, 2016; Aznar et al, 2015; Barbazan et al, 2016). What triggers such role-reversal, was unknown. We propose that when Wnt and growth factor pathways are deregulated concurrently during cancer progression [likely due to sequential mutations within each pathway, first in Wnt followed by growth factors], growth factor RTKs phosphorylate Daple’s PBM and augment the prometastatic Daple-dependent non-canonical Wnt signals, thereby fueling cancer dissemination.

In conclusion, we have provided evidence for how a novel signal transducer, Daple, provides a platform for complex crosstalk between non-canonical Wnt, growth factor RTKs, and G protein signaling cascades in multi-receptor driven diseases such as cancers. Findings also illuminate how such complex crosstalk between major signaling pathways coordinately shapes cellular phenotypes, sometimes even by reprogramming tumor suppressive pathways to aid tumor dissemination.

## FIGURE LEGENDS

**Figure 1-figure supplement 1:**
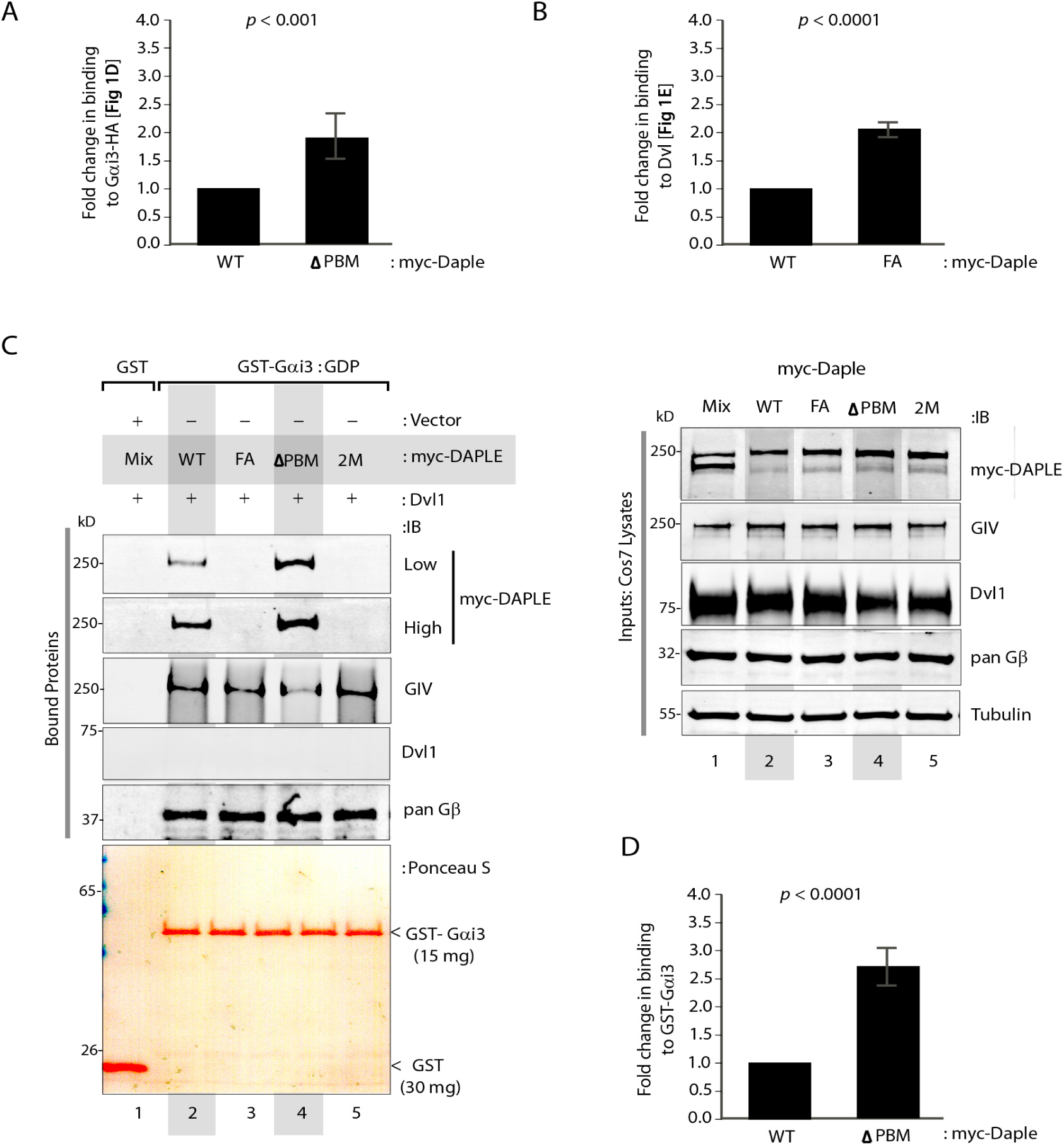
The <u>G</u>αi-<u>b</u>inding and <u>a</u>ctivating [GBA] and <u>P</u>DZ-<u>b</u>inding <u>m</u>otifs [PBM] in Daple allosterically inhibit each other’s functions. (**A**-**B**) Bar graphs display quantification of full-length myc-Daple bound to Gαi3-HA (Figure 1D) or Dvl (Figure 1E). Binding is expressed as fold change normalized to Daple-WT on the y-axis. Error bars represent mean ± S.D of three independent experiments. (**C**) Binding of full-length Daple to GST-Gαi3 is increased upon deletion of Daple PBM domain (ΔPBM). Equal aliquots of lysates of Cos7 cells expressing myc-Daple [WT and various mutants, as in Figure 1E] were used as the source of Daple in pulldown assays with bacterially expressed and purified GST-Gαi3 proteins immobilized on glutathione-Sepharose beads. Bound proteins were analyzed for Daple (myc), Gβ (positive control) and Dvl (negative control) by immunoblotting (IB). GIV, a paralogue of Daple that binds Gαi using a shared GBA motif, but does not contain a PBM domain and therefore, cannot bind Dvl was used as an internal control. As expected, GIV bound Gαi3 in all lysates tested. (**D**) Bar graphs display quantification of full-length myc-Daple bound to GST-Gαi3. Binding is expressed as fold change normalized to Daple-WT on y axis. Error bars represent mean ± S.D of three independent experiments.

**Figure 2-figure supplement 1:**
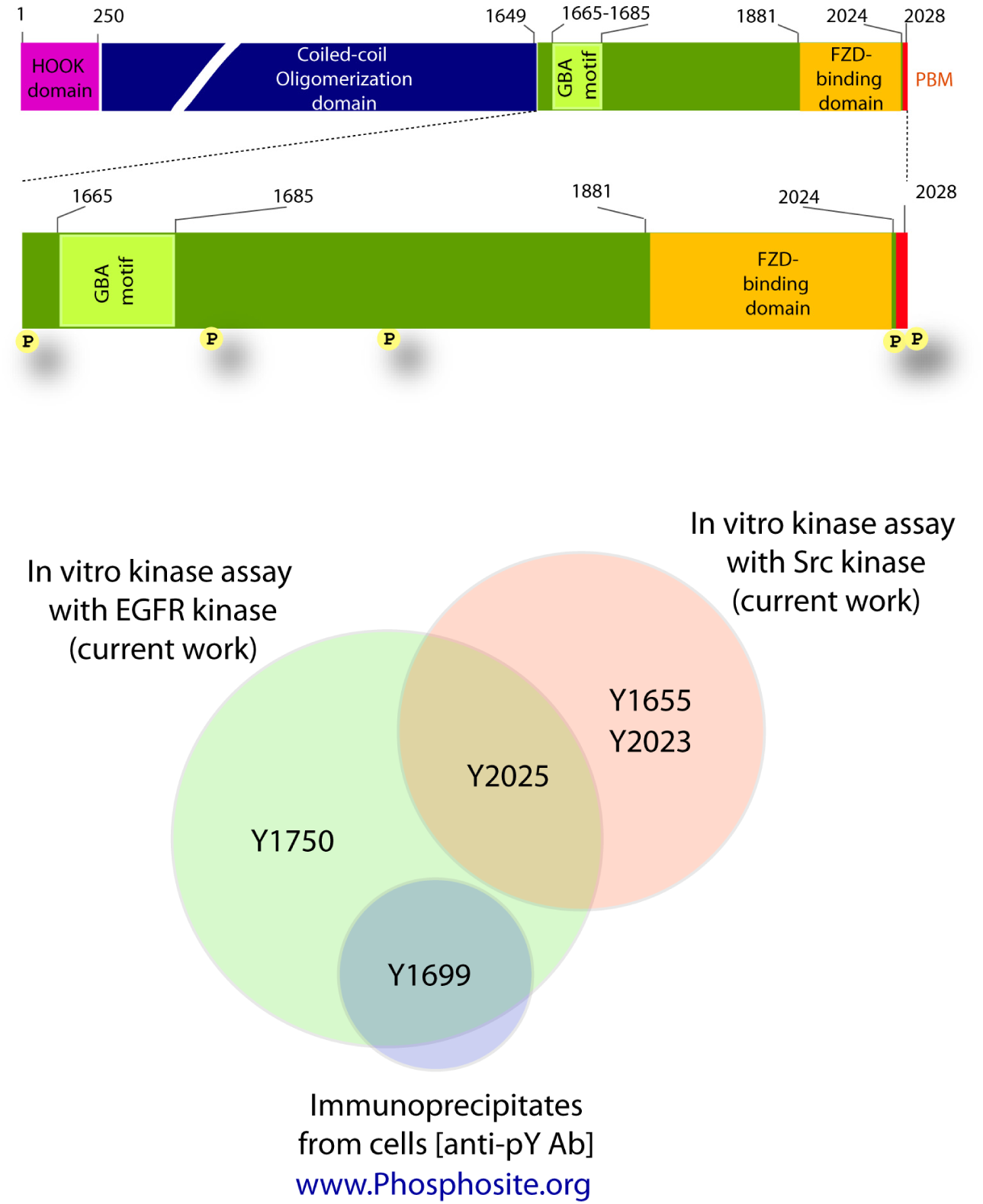
RTKs and non-RTKs target distinct sets of tyrosines within Daple’s C-terminus [aa 1650-2028]. *<u>Top</u>:* Schematic showing the distribution of 5 phosphorylated tyrosines on Daple’s C-terminus. *<u>Bottom</u>*: Venn diagram summarizing mass spectrometry findings. *In vitro* kinase assays carried out using EGFR and Src recombinant kinases [as in Figure 1C] were analyzed by mass spectrometry. His-Daple-CT protein used in these kinase assays contain 5 tyrosines. Of these, Y1655 and Y2023 were phosphorylated by Src, Y1699 and Y1750 by EGFR, and Y2025 was targeted by both kinases. Neither EGFR, nor Src phosphorylated Daple at Y1699, which is the only site reported to be phosphorylated in cells [www.phosphosite.org].

**Figure 2-figure supplement 2:**
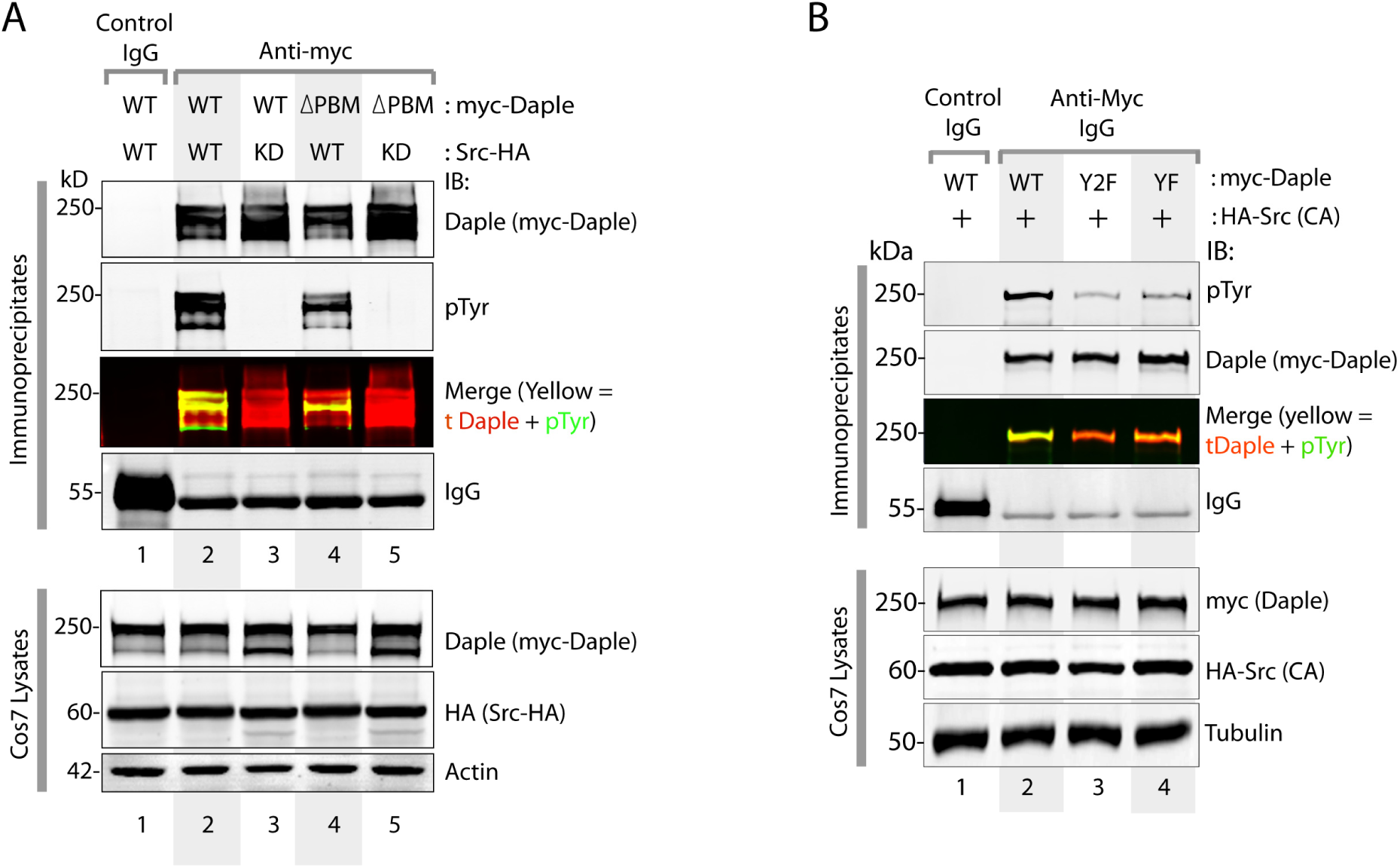
Src phosphorylates Daple at both Y2025 [which lies within the PBM] as well as Y2023 [that lies just outside the PBM]. (**A**) Daple was immunoprecipitated from equal aliquots of lysates of Cos7 cells co-expressing myc-Daple [WT or ΔPBM mutant] and HA-Src [WT or a kinase dead mutant (KD; K295R)] using anti-myc or control IgG. Immune complexes (top) were analyzed by dual color immunoblotting (IB) for total and tyrosine phosphorylated Daple using anti-Daple (red) and anti-pTyr mAb (green), respectively. Individual channels are shown in grayscale. Yellow pixels in merged pTyr (green) and Daple (red) panels show tyrosine phosphorylated Daple. Myc-Daple-WT was tyrosine phosphorylated in presence of Src-WT (lane 2) but not in presence Src-KD (lane 3). In the case of myc-Daple-ΔPBM (which lacks Y2025), tyrosine phosphorylation was detected in the presence of Src-WT, but not Src-KD (compare lanes 4 and 5), albeit to a lesser extent compared to myc-Daple-WT (compare lanes 2 and 4). These results indicate that Src phosphorylates Y2025 within Daple’s PBM, but also phosphorylates at additional sites outside that motif. (**B**) Daple was immunoprecipitated from equal aliquots of lysates of Cos7 cells co-expressing myc-Daple [WT, YF or Y2F mutant] and Src-HA [constitutively active Y527F mutant; CA] using anti-myc or control IgG. Immune complexes were analyzed by dual color immunoblotting (IB) for total and tyrosine phosphorylated Daple using anti-Daple (red) and anti-pTyr mAb (green), respectively as in **A**. Yellow pixels in merged pTyr (green) and Daple (red) panels show that tyrosine phosphorylation of Daple was maximal in Daple-WT and reduced significantly (~by 80%) in Daple-Y2F, which lacks both Y2025 and Y2023 (compare lanes 2 and 3). Tyrosine phosphorylation of Daple-YF was intermediate between Daple-WT and Daple-Y2F (compare lanes 2-4). These findings indicate that Src phosphorylates Daple at both Y2025 and 2023. The residual phosphorylation observed in Daple-Y2F (lane 3) could be the third Src site identified by mass spectrometry analysis (see Figure2-figure supplement 1).

**Figure 2-figure supplement 3:**
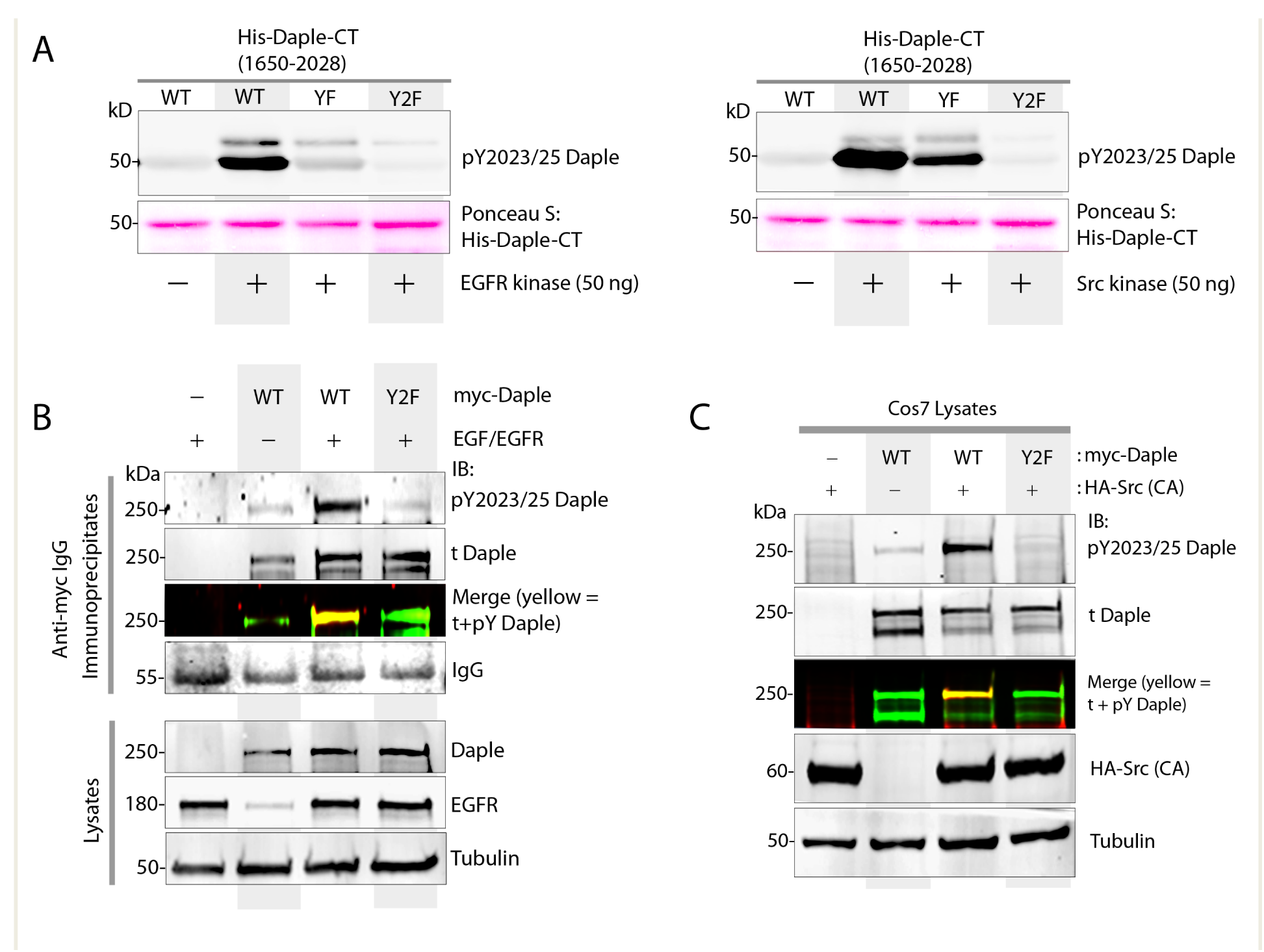
Validation of the phospho-specific anti-pY2023/25 Daple antibody. (**A**) *In vitro* kinase assays were carried out using either EGFR or Src recombinant kinase and purified His-Daple-CT [WT or non-phosphorylatable mutants Y2025F (YF) and Y2023/2025F (Y2F)] exactly as in Figure 2B. Entire reactions were separated by SDS PAGE and analyzed by ponceau S staining [to confirm equal loading of substrate proteins] and immunoblotting for tyrosine phosphorylated Daple using anti-pY2023/25 Daple antibody. (**B, C**) Lysates of Cos7 cells co-expressing myc-Daple [WT or Y2F mutant] and EGFR (**B**) or constitutively active HA-Src mutant (Y527F; CA) (**C**) were analyzed by dual color immunoblotting for total and tyrosine phosphorylated Daple using myc (green; t Daple) and anti-pY2023/25 Daple (red) antibodies, respectively. Individual channels are shown in grayscale. Yellow pixels in merged t Daple (green) and pY2023/25 Daple (red) panels show that the anti-pY2023/25 Daple antibody detects tyrosine phosphorylated Daple exclusively when the two phosphorylatable tyrosines are intact (i.e., in Daple-WT) but not when the two tyrosines are mutated to Phe (i.e., in Daple-Y2F). The anti-pY2023/25 Daple antibody does not detect unphosphorylated Daple-WT, or other phosphosites on Daple-Y2F that are phosphorylated by EGFR and Src (see Figure2-figure supplement 1).

**Figure 3-figure supplement 1:**
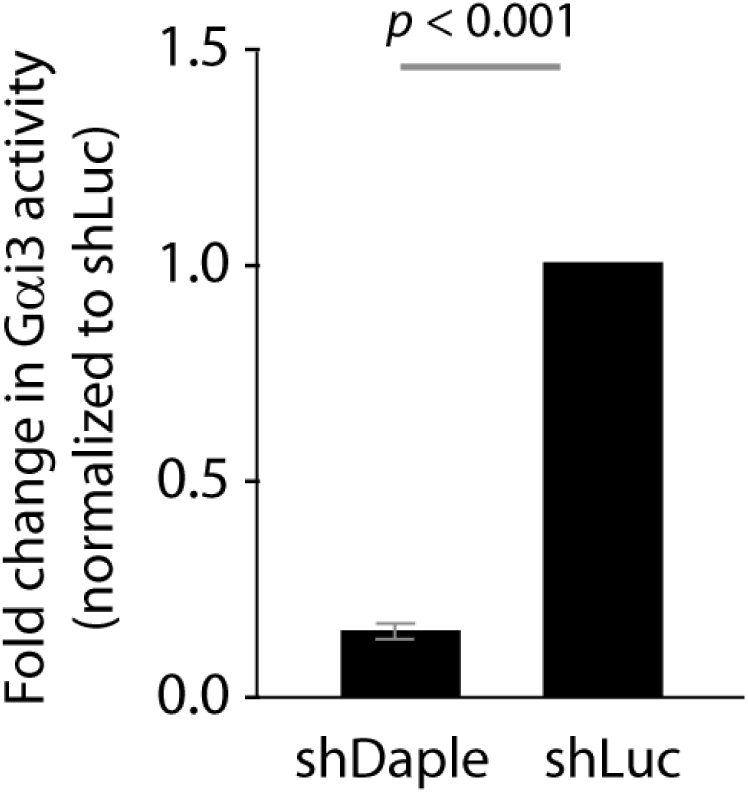
Daple is required for activation of Gαi3 downstream of EGF/EGFR. Bar graphs display fold change in Gαi3 activity observed in Figure 3A. Error bars represent mean ± S.D of three independent experiments.

**Figure 4-figure supplement 1:**
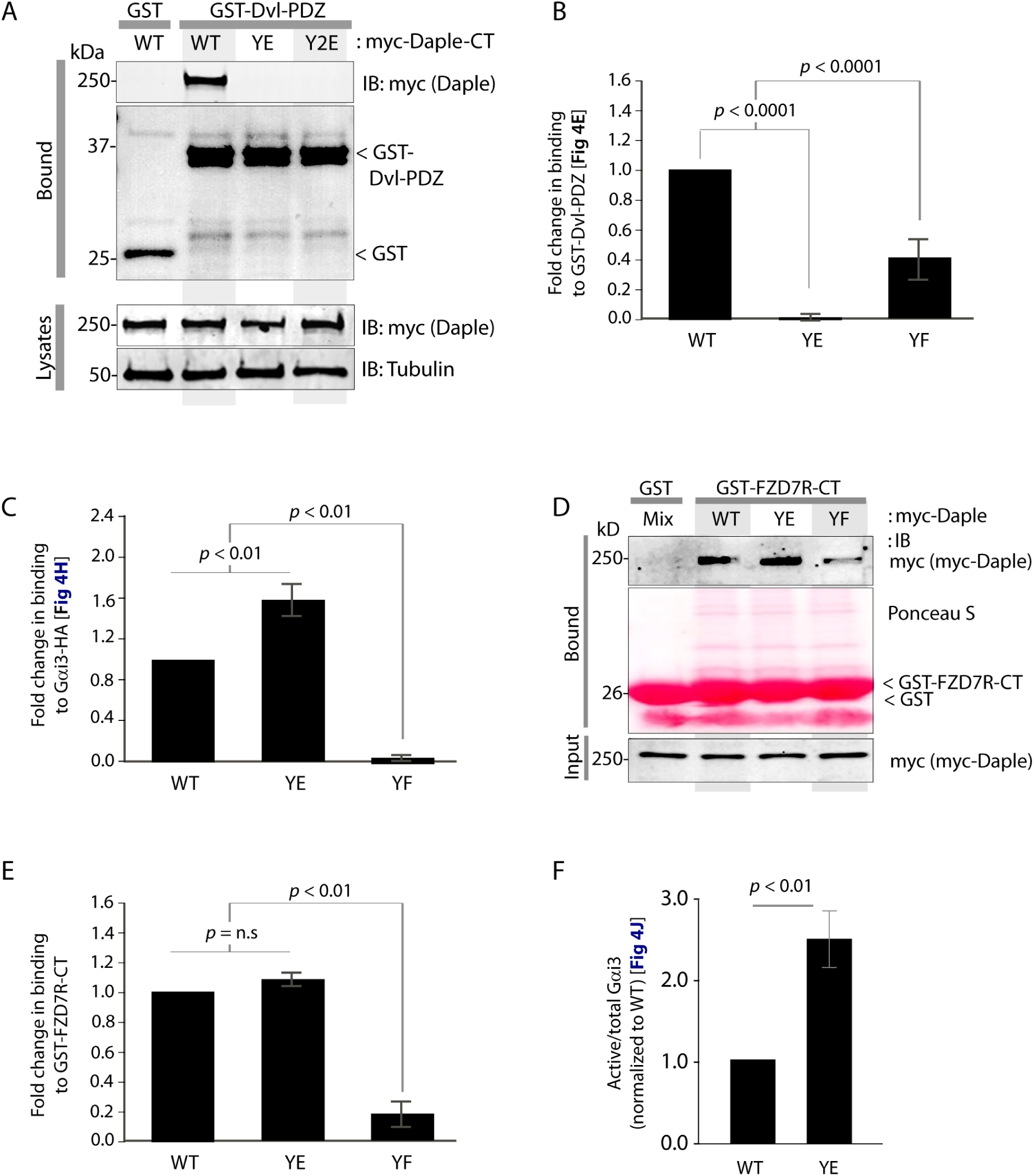
Phosphorylation of Y2025 within Daple’s PBM (mimicked by Y→E mutation) inhibits its ability to bind Dvl, enhances binding to Gαi, and has no impact on binding to FZD7R. **(A)** Lysates of Cos7 cells expressing myc-Daple WT, Y2025E (YE) or Y2023, 2025E (Y2E) were incubated in presence of GST-Dvl2-PDZ. Bound complexes were analyzed for Daple using anti-myc mAb by immunoblotting (IB). As expected, Daple-WT bound GST-Dvl2-PDZ. No binding was detected in the case of Daple-YE or Y2E, indicating that phosphorylation on Y2025 (a common target for both RTKs and non-RTKs) within Daple’s PBM may be sufficient to maximally disrupt Daple:Dvl complexes. (**B**) Bar graphs display fold change in binding between GST-Dvl-PDZ and Daple (WT and mutants) observed in Figure 4E. Error bars represent mean ± S.D of three independent experiments. (**C**) Bar graphs display fold change in binding between Gαi3-HA and Daple (WT and mutants) observed in Figure 4H. Error bars represent mean ± S.D of three independent experiments. (**D, E**) Phosphorylation-mimic mutant of Y2025 doesn’t affect binding of Daple to the cytoplasmic tail of Frizzled 7 receptor (FZD7R). Lysates of Cos7 cells expressing myc-Daple WT, YE or YF were incubated in presence of GST-FZD7R-CT. Bound complexes were analyzed for Daple using anti-myc mAb by immunoblotting (IB). Daple-WT and Daple-YE bound similarly to GST-FZD7R-CT. Compared to Daple-WT or Daple-YE, Daple-YF bound to a lesser extent (**D**). Bar graphs in **E** display fold change in binding between GST-FZD7R-CT and Daple (WT and mutants) observed in **D**. Error bars represent mean ± S.D of three independent experiments. (**F**) Bar graphs display Gαi3 activity observed in Figure 4J. Error bars represent mean ± S.D of three independent experiments.

**Figure 5-figure supplement 1:**
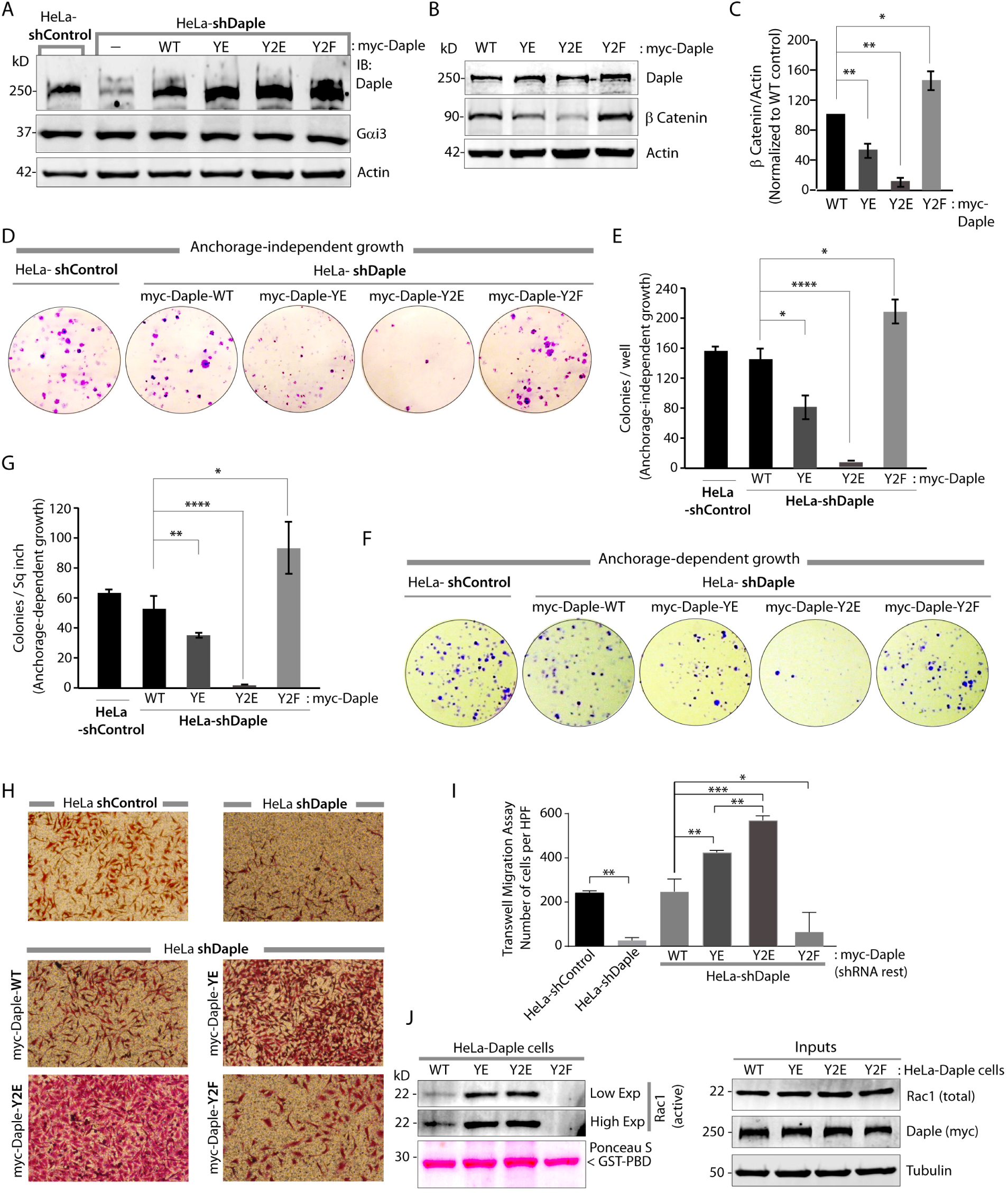
Tyrosine-phosphorylation of Daple’s PBM is essential for inhibition of colony growth and enhancement of cell migration. (**A**) Generation of HeLa cell lines stably expressing various Daple constructs (see also *Methods).* Whole-cell lysates from control (shControl) or Daple-depleted (shDaple) HeLa cells stably expressing Daple-WT, YE, Y2E and Y2F were analyzed for Daple, Gαi3 and actin by immunoblotting (IB). Band densitometry (using LiCOR Odyssey) confirmed that various Daple constructs were expressed at levels ~1.0-1.5 fold of endogenous. (**B, C**) Constitutive phosphorylation of Daple on Y2023 and Y2025 [mimicked by expression of Daple-Y2E mutant] decreases the levels of β-catenin; expression of a non-phosphorylatable Y2F mutant has an opposite effect. Equal aliquots of lysates of various HeLa Daple cell lines were analyzed for β-catenin by immunoblotting (IB; **B**). Bar graphs in **C** display quantification of β-catenin in **A**. Error bars represent mean ± S.D of three independent experiments. (**D-G**) Constitutive phosphorylation of Daple on Y2023 and Y2025 [mimicked by expression of Daple-Y2E mutant] inhibits both anchorage-independent (**D, E**) and anchorage-dependent (**F, G**) colony growth; expression of a nonphosphorylatable Y2F mutant has an opposite effect. Various HeLa Daple cells lines were analyzed for their ability to form colonies in soft agar or on plastic for 2–3 weeks (see *Methods*). Panel **D** shows the photograph of the crystal violet-stained wells of a 6-well plate. The number of colonies was counted by light microscopy throughout the depth of the matrix in 15 randomly chosen fields. In panel **E**, bar graphs display the number of colonies (y axis) seen for each cell line in **D**. Panel **F** shows the photograph of the crystal violet-stained wells of a 6-well plate. The number of colonies was counted by ImageJ (Colony counter). Panel **G** shows bar graphs that display the % inhibition of colony formation (y axis) seen in each condition normalized to shControl HeLa cells. (**H, I**) Constitutive phosphorylation of Daple on Y2023 and Y2025 [mimicked by expression of Daple-Y2E mutant] enhances cell migration; expression of a non-phosphorylatable Y2F mutant has an opposite effect. Various HeLa Daple cells lines were analyzed for their ability to migrate in transwell assays (see *Methods*). Panel **H** shows the photograph of cells after 24 hr of migration, fixed and stained with Giemsa. Bar graphs in **I** display the number of migrating cells from 20 high power fields (HPFs) per experiment. Data are presented as mean ± S.D; n = 3. (**J**) Constitutive phosphorylation of Daple on Y2023 and Y2025 [mimicked by expression of Daple-Y2E mutant] enhances Rac1 activity; expression of a nonphosphorylatable Y2F mutant has an opposite effect. HeLa Daple cell lines grown in 10% serum were analyzed for Rac1 activation by pulldown assays using GST-PBD (see *Methods*). Bound proteins (active GTP-bound Rac1) were analyzed by immunoblotting (IB)

**Figure 5-figure supplement 2:**
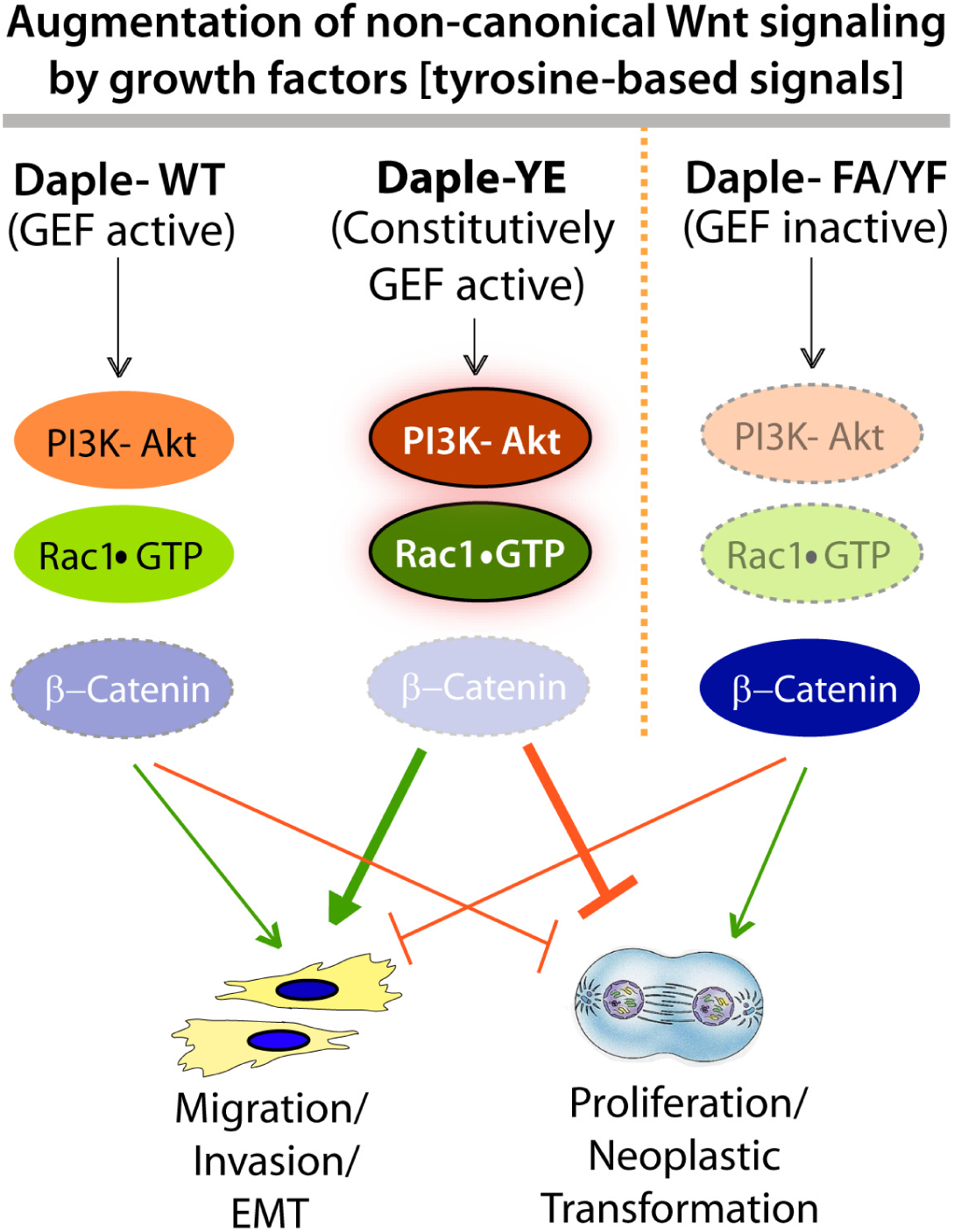
Schematic summarizing the signaling profiles and phenotypes triggered by various Daple mutants used in the current work. We previously showed that Daple-WT antagonizes canonical β-catenin/Wnt signaling, but enhances Rac1 and Akt signals; consequently, proliferation is suppressed but migration/invasion and EMT is triggered (Aznar et al, 2015). A GEF-deficient mutant (Daple-F1675A; FA), which cannot bind or activate Gαi showed an opposite phenotype. Here we show that constitutive phosphorylation of Daple [as mimicked by Daple-Y2025E] further enhances the signaling programs initiated by Daple-WT. Expression of a non-phosphorylatable YF mutant which, like Daple-FA, cannot bind or activate Gαi recapitulated signaling programs and phenotypes seen in cells expressing Daple-FA.

**Figure 6-figure supplement 1:**
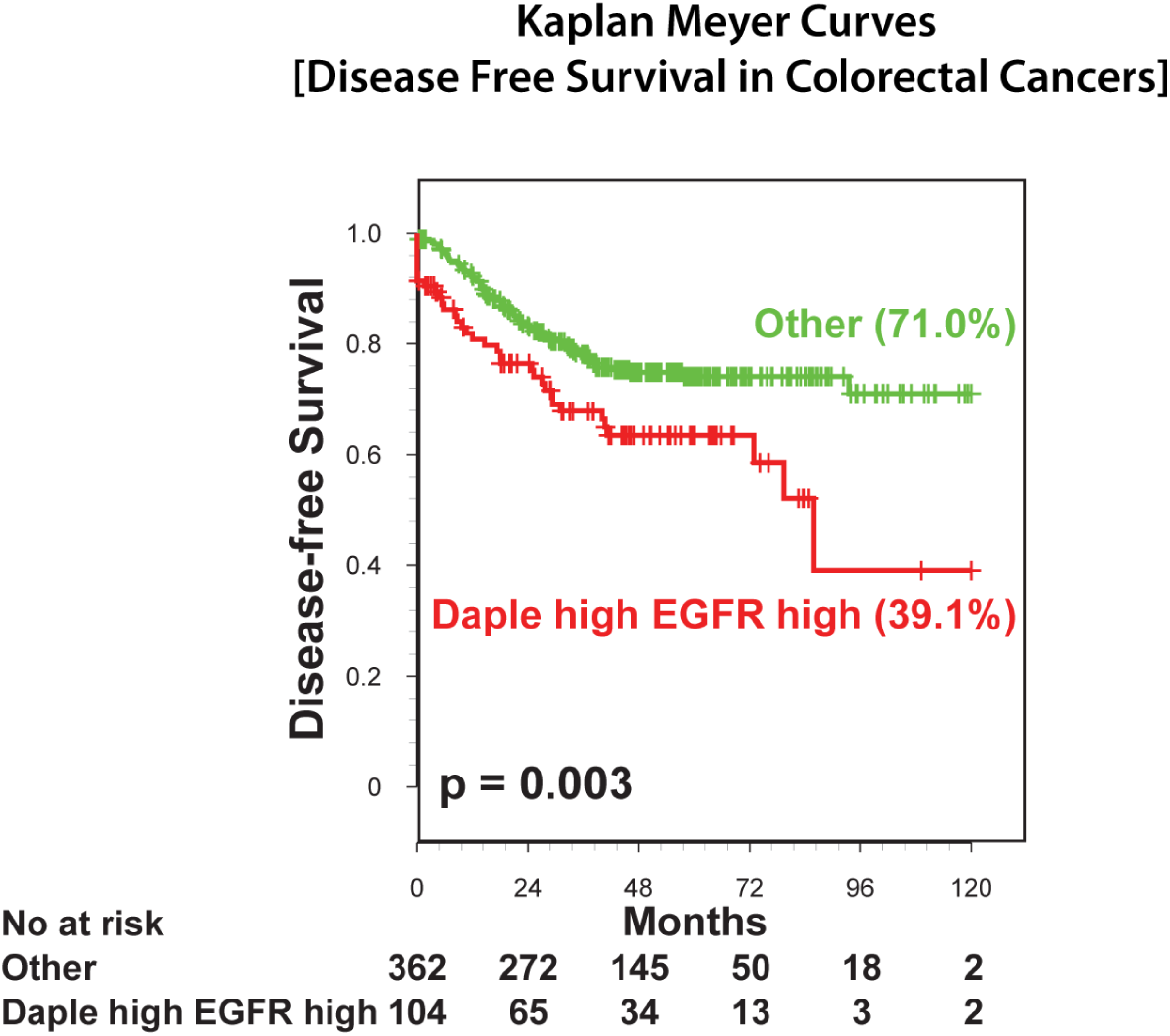
Concurrent high Daple (CCDC88C) and EGFR mRNA levels in colon cancers constitutes a unique poor prognostic signature with lower disease-free survival (DFS) compared to all other tumors. 466 patients with colon cancer were stratified into high vs low *Daple* (CCDC88C) and *EGFR* expression levels using Hegemon software (see *Methods*; Figure 6A). DFS analysis using Kaplan-Meier curves showed that concurrent high expression levels for EGFR and Daple carried a significantly worse prognosis when compared to all other subgroups [‘Others’] combined [high EGFR/ low Daple, low EGFR/low Daple, and low EGFR/high Daple].

## Materials and Methods

### Reagents and Antibodies—

Unless otherwise indicated, all reagents were of analytical grade and obtained from Sigma-Aldrich (St. Louis, MO). Cell culture media were purchased from Invitrogen (Grand Island, NY). All restriction endonucleases and Escherichia coli strain DH5α were purchased from New England Biolabs (Ipswich, MA). E. coli strain BL21 (DE3), phalloidin-Texas Red were purchased from Invitrogen. Genejuice transfection reagent was from Novagen (Madison, WI). PfuUltra DNA polymerase was purchased from Stratagene (La Jolla, CA). Goat anti-rabbit and goat anti-mouse Alexa Fluor 680 or IRDye 800 F(ab′)2 used for immunoblotting were from Li-Cor Biosciences (Lincoln, NE). Mouse anti-His, anti-α tubulin and anti-actin were obtained from Sigma; anti-Myc and anti-HA were obtained from Cell Signaling Technology (Beverly, MA) and Covance (Princeton, NJ), respectively. Mouse monoclonal antibody against phosphotyrosine (pTyr, Cat # 610000) was obtained from BD Transduction Laboratories (San Jose, CA). Rabbit anti–pan-Gβ (M-14), anti-Gαi3, anti-Dvl and anti-β-catenin were obtained from Santa Cruz Biotechnology (Dallas, TX); antiphospho-Akt (S473) and anti-pan Akt were obtained from Cell Signaling Technology; anti-Rac1 was obtained from BD Transduction Laboratories. Rabbit polyclonal anti-Daple (total) antibody were generated in collaboration with Millipore (Carlsbad, CA) using the C-terminus of Daple (aa 1660-2028) as an immunogen and validated previously (Aznar et al, 2015).

An affinity purified rabbit polyclonal phospho-specific anti-Daple (pY2023, pY2025) antibody was generated using a phosphopeptide corresponding to the PBM motif in human Daple as immunogen by 21^st^ Century Biochemicals (Marlboro, MA). Briefly, rabbits were immunized with a 1:1 mix of 2 phosphopeptides-- Ahx-PQTVWYE[pY]GCV-OH and Ahx-PQTVW[pY]E[pY]GCV-OH, and purified by adsorbing antibodies against the non-phosphorylated sequence using a Ahx-PQTVWYEYGCV-OH peptide.

### Plasmid Constructs and Mutagenesis—

Cloning of N-terminally tagged myc-Daple was carried out as described previously (Aznar et al, 2015). All subsequent site-directed mutagenesis and truncated constructs (myc-Daple full length F1675A (FA), myc-Daple deleted from aa 2025-2028 (ΔPBM), myc-Daple FA+ΔPBM (2M), myc-Daple Y2025E (YE), Y2023-2025E (Y2E), Y2025F (YF), and Y2023-2025F (Y2F) were carried out using myc-Daple-WT as template using Quick Change as per manufacturer’s protocol. The His-Daple-CT WT, YF and Y2F constructs (1650-2028 aa) used for in vitro protein-protein interaction assays were cloned from myc-Daple pcDNA 3.1 and inserted within the pGEX-4T vector, respectively, between *NdeI*/*EcoRI* restriction sites.

Cloning of rat Gα-proteins into pGEX-4T-1 (GST-Gαi3 and Gαi3-HA) have been described previously (Garcia-Marcos et al, 2010; Garcia-Marcos et al, 2009; Garcia-Marcos et al, 2011c; Ghosh et al, 2010; Ghosh et al, 2008). C-terminal HA-tagged c-Src for mammalian expression was generated by cloning the entire coding sequence into pcDNA 3.1 between XhoI and EcoRI. HA-Src Y527F (constitutively active; CA) and HA-Src K295R (kinase dead; KD) were generated by site-directed mutagenesis using a Quick Change kit (Stratagene, San Diego, CA) as per manufacturer’s protocol. All HA-Src constructs have been previously validated by us (Lin et al, 2011). Several constructs used in this work were generous gifts from other investigators: Un-tagged-EGFR was from Marilyn G. Farquhar (UCSD, La Jolla,CA) (Ghosh et al, 2010); mouse Dvl1 was from Mikhail V. Semenov (Harvard Medical School, Boston, MA); GST-Dvl2-PDZ was from Raymond Habas (Temple University, Philadelphia, PA); GST-tagged FZD7-CT construct (Yao et al, 2004) was from Ryoji Yao (Japanese Foundation of Cancer Research institute, Japan); GST-PBD (PDZ-Binding Domain) was from Gary Bokoch (The Scripps Research Institute, La Jolla, CA). All these constructs have been previously used and validated by us in our recently published work on Daple (Aznar et al, 2015).

Daple shRNA constructs have been validated and are described in detail elsewhere (Aznar et al, 2015).

### Protein Expression and Purification—

GST and His-tagged recombinant proteins were expressed in E. coli strain BL21 (DE3) (Invitrogen) and purified as described previously (Garcia-Marcos et al, 2011a; Ghosh et al, 2010; Ghosh et al, 2008). Briefly, bacterial cultures were induced overnight at 25 °C with 1 mM isopropylβ-D-1-thiogalactopyranoside (IPTG). Pelleted bacteria from 1 L of culture were resuspended in 20 mL GST-lysis buffer [25 mM Tris·HCl, pH 7.5, 20 mM NaCl, 1 mM EDTA, 20% (vol/vol) glycerol, 1% (vol/vol) Triton X-100, 2×protease inhibitor mixture (Complete EDTA-free; Roche Diagnostics)] or in 20mL His-lysis buffer [50 mM NaH2PO4 (pH 7.4), 300 mM NaCl, 10 mM imidazole, 1% (vol/vol)Triton X100, 2×protease inhibitor mixture (Complete EDTA-free; Roche Diagnostics)] for GST or His-fused proteins, respectively. After sonication (three cycles, with pulses lasting 30 s/cycle, and with 2 min interval between cycles to prevent heating), lysates were centrifuged at 12,000×gat 4 °C for 20 min. Except for GST-FZD7-CT and GST-PBD constructs (see *in vitro* GST pulldown assay section), solubilized proteins were affinity purified on glutathione-Sepharose 4B beads (GE Healthcare), dialyzed overnight against PBS, and stored at−80 °C.

### Transfection; Generation of Stable Cell Lines and Cell Lysis—

Transfection was carried out using either PEI as previously described (Longo et al, 2013) or Genejuice from Novagen for DNA plasmids following the manufacturers’ protocols. HeLa cell lines stably expressing Daple constructs were selected after transfection and in the presence of Dulbecco’s Modified Eagle’s medium (DMEM) supplemented with 800 μg/ml G418 for 6 weeks. The resultant multiclonal pool was subsequently maintained in the presence of 500 μg/ml G418. Unless otherwise indicated, for assays involving serum starvation, serum concentration was reduced to 0.2% Fetal Bovine Serum (FBS) overnight for HeLa cells and 0% FBS for Cos7.

Whole-cell lysates were prepared after washing cells with PBS prior to resuspending and boiling them in sample buffer. Cold PBS [4°C] was used whenever ligand stimulation was carried out for designated time periods. Lysates used as a source of proteins in immunoprecipitation or pull-down assays were prepared by resuspending cells in Tx-100 lysis buffer [20 mM HEPES, pH 7.2, 5 mM Mg-acetate, 125 mM K-acetate, 0.4% Triton X-100, 1 mM DTT, supplemented with sodium orthovanadate (500 μM), phosphatase from Sigma-Aldrich and protease from Roche Life Science (Indianapolis, IN) inhibitor cocktails], after which they were passed through a 28G needle at 4°C, and cleared by centrifuging (10,000 x g for 10 min) before use in subsequent experiments.

### Quantitative Immunoblotting—

For immunoblotting, protein samples were separated by SDS-PAGE and transferred to PVDF membranes (Millipore). Membranes were blocked with PBS supplemented with 5% nonfat milk (or with 5% BSA when probing for phosphorylated proteins) before incubation with primary antibodies. Infrared imaging with two-color detection and band densitometry quantifications were performed using a Li-Cor Odyssey imaging system exactly as done previously (Aznar et al, 2015). All Odyssey images were processed using Image J software (NIH) and assembled into figure panels using Photoshop and Illustrator software (Adobe).

### In vitro GST pulldown and Immunoprecipitation Assays –

Purified GST-Gαi3, GST-DVL-PDZ or GST alone (5 μg) were immobilized on glutathione-Sepharose beads and incubated with binding buffer [50 mM Tris-HCl (pH 7.4), 100 mM NaCl, 0.4% (v:v) Nonidet P-40, 10 mM MgCl_2_, 5 mM EDTA, 30 μM GDP, 2 mM DTT, protease inhibitor mixture] for 90 min at room temperature as described before (Garcia-Marcos et al, 2011a; Ghosh et al, 2010; Ghosh et al, 2008; Lin et al, 2011). Lysates (~250 μg) of Cos7 cells expressing appropriate myc-Daple constructs or purified His-Daple-CT (aa 1650-2028) protein (3 μg) were added to each tube, and binding reactions were carried out for 4 h at 4°C with constant tumbling in binding buffer [50 mM Tris-HCl (pH 7.4), 100 mM NaCl, 0.4% (v:v) Nonidet P-40, 10 mM MgCl_2_, 5 mM EDTA, 30 μM GDP, 2 mM DTT]. Beads were washed (4x) with 1 mL of wash buffer [4.3 mM Na_2_HPO_4_, 1.4 mM KH_2_PO_4_ (pH 7.4), 137 mM NaCl, 2.7 mM KCl, 0.1 % (v:v) Tween 20, 10 mM MgCl_2_, 5 mM EDTA, 30 μM GDP, 2 mM DTT] and boiled in Laemmli’s sample buffer. Immunoblot quantification was performed by infrared imaging following the manufacturer’s protocols using an Odyssey imaging system (Li-Cor Biosciences).

GST-FZD7-CT and GST-PBD constructs were immobilized on glutathione-Sepharose beads directly from bacterial lysates by overnight incubation at 4°C with constant tumbling. Next morning, GST-FZD7-CT or GST-PBD immobilized on glutathione beads were washed and subsequently incubated with cell lysates at 4°C with constant tumbling. Washes and immunoblotting were performed as previously (Aznar et al, 2015).

For immunoprecipitation, cell lysates (~1-2 mg of protein) were incubated for 4 h at 4°C with 2 μg of appropriate antibody, anti-HA mAb (Covance) for HA-Gαi3, anti-myc (from Cell Signaling) mAb for myc-Daple, anti-Dvl mAb (from Santa Cruz Biotechnology) or their respective pre-immune control IgGs. Protein G (for all mAbs) Sepharose beads (GE Healthcare) were added and incubated at 4°C for an additional 60 min. Beads were washed in PBS-T buffer [4.3 mM Na_2_HPO_4_, 1.4 mM KH_2_PO_4_, pH 7.4, 137 mM NaCl, 2.7 mM KCl, 0.1 % (v:v) Tween 20, 10 mM MgCl_2_, 5 mM EDTA, 2 mM DTT, 0.5 mM sodium orthovanadate] and bound proteins were eluted by boiling in Laemmli’s sample buffer.

### In vitro and in-cellulo Kinase Assays—

In vitro kinase assays were performed using bacterially expressed His (6 x His, hexahistidine) tagged Daple-CT (His-Daple-CT, aa 1650-2028) proteins (~5 μg per reaction), and ~50 ng recombinant kinases which were obtained commercially (SignalChem, Canada). Reactions were started by adding ~1000 μM ATP and carried out at 25°C for 30 min in tyrosine kinase buffer [60 mM HEPES pH 7.5, 5 mM MgCl_2_, 5 mM MnCl_2_, 3 μM sodium orthovanadate]. Reactions were stopped by addition of Laemmli sample buffer and boiling at 100°C.

For cellular phosphorylation assays using overexpressed Daple, Myc-tagged Daple was coexpressed with untagged EGFR or HA-tagged Src-WT, HA-tagged Src-kinase-dead [KD] or HA-tagged Src-constitutively active [CA]. To determine if EGFR phosphorylates Daple, cells were serum starved for 12-16 h, and stimulated with EGF (50 nM) for 10 min. Cells were preincubated for 1 h with Src inhibitor, PP2 [Calbiochem] as done previously (Lin et al., 2011). Reactions were stopped using PBS chilled at 4°C, supplemented with 200 μM sodium orthovanadate, and immediately scraped and lysed for immunoprecipitation.

Phosphoproteins in both *in vitro* and *in cellulo* assays were visualized by immunoblotting with either a pan-pTyr antibody [BD Biosciences] or a phosphospecific anti-pTyr 2023-2025-Daple rabbit polyclonal antibody [21^st^ Century Biochemicals].

### Mass Spectrometry—

These studies were carried out as described previously (Guttman et al, 2009; McCormack et al, 1997). The key steps are summarized below.

Sample preparation: His-Daple-CT proteins used in various *in vitro* kinase assays were diluted in TNE (50 mM Tris, pH 8.0, 100 mM NaCl, 1 mM EDTA) buffer. RapiGest SF reagent (Waters Corp.) was added to the mix to a final concentration of 0.1% and samples were boiled for 5 min. TCEP (Tris (2-carboxyethyl) phosphine) was added to 1 mM (final concentration) and the samples were incubated at 37°C for 30 min. Subsequently, the samples were carboxymethylated with 0.5 mg/ml of iodoacetamide for 30 min at 37°C followed by neutralization with 2 mM TCEP (final concentration). Proteins samples prepared as above were digested with trypsin (trypsin:protein ratio - 1:50) overnight at 37°C. RapiGest was degraded and removed by treating the samples with 250 mM HCl at 37°C for 1 h followed by centrifugation at 14000 rpm for 30 min at 4°C. The soluble fraction was then added to a new tube and the peptides were extracted and desalted using C18 desalting columns (Thermo Scientific, PI-87782).

LC-MS-MS: Trypsin-digested peptides were analyzed by ultra high pressure liquid chromatography (UPLC) coupled with tandem mass spectroscopy (LC-MS/MS) using nano-spray ionization. The nano-spray ionization experiments were performed using a TripleTof 5600 hybrid mass spectrometer (ABSCIEX) interfaced with nano-scale reversed-phase UPLC (Waters corporation nano ACQUITY) using a 20 cm-75-micron ID glass capillary packed with 2.5-μm C18 (130) CSH™ beads (Waters corporation). Peptides were eluted from the C18 column into the mass spectrometer using a linear gradient (5–80%) of ACN (Acetonitrile) at a flow rate of 250 μl/min for 1h. The buffers used to create the ACN gradient were: Buffer A (98% H_2_O, 2% ACN, 0.1% formic acid, and 0.005% TFA) and Buffer B (100% ACN, 0.1 % formic acid, and 0.005% TFA). MS/MS data were acquired in a data-dependent way the MS1 data was acquired for 250 ms at m/z of 400 to 1250 Da and the MS/MS data was acquired from m/z of 50 to 2,000 Da. The Independent data acquisition (IDA) parameters were as follows; MS1-TOF acquisition time of 250 milliseconds, followed by 50 MS2 events of 48 milliseconds acquisition time for each event. The threshold to trigger MS2 event was set to 150 counts when the ion had the charge state +2, +3 and +4. The ion exclusion time was set to 4 seconds. Finally, the collected data were analyzed using Protein Pilot 4.5 (ABSCIEX) for peptide identifications.

### Gαi activity as determined by conformational anti-Gαi•GTP mAb—

These assays were carried out exactly as done previously (Aznar et al, 2015; Lin et al, 2014). Cells were maintained overnight at steady-state in a media containing 0.2% FBS [if an EGF stimulation was performed] or maintained at steady-state in media containing 10% FBS prior to lysis. For immunoprecipitation of active Gαi3, freshly prepared cell lysates (2-4 mg) were incubated for 30 min at 4°C with the conformational Gαi:GTP mouse antibody (1 μg) (Lane et al, 2008) or with control mouse IgG. Protein G Sepharose beads from GE Healthcare (Pittsburgh, PA) were added and incubated at 4°C for additional 30 min (total duration of assay is 1 h). Beads were immediately washed 3 times using 1 ml of lysis buffer (composition exactly as above; no nucleotides added) and immune complexes were eluted by boiling in SDS as previously described (Aznar et al, 2015; Lin et al, 2014).

### Measurement of Rac1 activity—

These assays were carried out exactly as done previously (Aznar et al, 2015) with slight modifications in FBS concentration. Briefly, to analyze the role of phosphotyrosine Daple in the regulation of Rac1 activity we used Daple-depleted HeLa cell lines stably expressing Daple-WT or tyrosine mutants. Cells were maintained overnight at steady-state in a media containing 10% FBS prior to lysis. Lysis was carried out first in RIPA buffer [20 mM HEPES pH 7.4, 180 mM NaCl, 1 mM EDTA, 1% Triton X-100, 0.5% sodium deoxycholate, 0.1% SDS, supplemented with 1mM DTT, sodium orthovanadate (500 μM), phosphatase (Sigma), and pro-tease (Roche) inhibitor mixtures] for 15 min on ice, and then for an additional 15 min after addition of an equal volume of Triton X-100 lysis buffer [20 mM Hepes (pH 7.2), 5 mM Mg-acetate, 125 mM K- acetate, 0.4% Triton X-100, 1 mM DTT, supplemented with sodium orthovanadate (500 μM), phosphatase (Sigma), and protease (Roche) inhibitor mixtures]. During the second 15 min of incubation, cells were broken by passing through a 28-gauge needle at 4 °C and lysates were subsequently cleared (10,000×g for 10 min) before use. Equal aliquots of lysates were incubated with bead-bound GST-PBD for 1h at 4°C with constant tumbling. Beads were washed in PBS-T buffer [4.3 mM Na_2_HPO_4_, 1.4 mM KH_2_PO_4_, pH 7.4, 137 mM NaCl, 2.7 mM KCl, 0.1% (v:v) Tween 20, 10 mM MgCl_2_, 5 mM EDTA, 2 mM DTT, 0.5 mM sodium orthovanadate] and bound proteins were eluted by boiling in Laemmli’s sample buffer.

### Transwell Cell Migration—

Chemotactic cell migration assays were performed using Corning Transwell plates according to the manufacturer’s protocol exactly as done previously (Aznar et al, 2015) with slight modifications in FBS concentration. HeLa cells were trypsinized, counted, and placed in a Transwell with media containing no FBS (75000 cells/well). Media in the bottom chamber of each well were supplemented with 10% FBS to trigger chemotactic migration. Cells were allowed to migrate for 24 hr and fixed prior to staining. Cells that had successfully migrated to the side of the permeable membrane facing the bottom chamber were visualized by staining the membrane with crystal violet. Cell migration (expressed as number of cells/high-power field) was quantified by analyzing 15–20 random fields per membrane insert per condition for the number of Giemsa stained cells. Each experiment was repeated 3 times [biological repeats]; 3 technical repeats were included during each biological repeat.

### 3D modeling of Dvl2-PDZ domain bound to the PDZ-binding motif (PBM) of Daple—

The approximate model of the complex was built using the ICM homology modeling platform (Cardozo et al, 1995). Structures of Dvl2-PDZ domain co-crystallized with various peptides were collected from the Pocketome (Kufareva et al, 2012). A structural alignment of the bound peptides was built, and the PDM of Daple was aligned onto it, suggesting the highest sequence homology with the C1 peptide [PDB 3cbx (Zhang et al, 2009)]. The initial model of the DVL2-PDZ: Daple-PBM complex was built by assigning the backbone coordinates of both target molecules to their counterparts in the template (PDB 3cbx). The model was further refined via conformational sampling of peptide and PDB domain residue side chains in internal coordinates, followed by full-atom local backbone minimization in the presence of harmonic distance restraints maintaining the secondary structure of the complex.

### Anchorage-dependent tumor growth assay—

Anchorage-dependent growth was monitored on solid (plastic) surface as performed by us previously (Aznar et al, 2015) with slight modifications in FBS concentration. Briefly, approximately ~1000 HeLa cells stably expressing various Daple constructs were plated in 6-well plates and incubated in 5% CO_2_ at 37°C for ~2 weeks in 10% FBS growth media. Colonies were then stained with 0.005% crystal violet for 1 h. Each experiment was repeated 3 times [biological repeats]; 3 technical repeats were included during each biological repeat.

### Anchorage-independent tumor growth assay--

Anchorage-independent growth of HeLa cells was analyzed in agar as performed by us previously (Aznar et al, 2015) with slight modifications in FBS concentration. Briefly, petri plates (60 mm) were pre-layered with 3 ml 1% Bacto agar (Life Technologies) in DMEM containing 10% FBS. Approximately ~5000 HeLa cells stably expressing various Daple constructs were then plated on top in 3 ml of 0.3% agar–DMEM with 10% FBS. All assays were carried out using three replicate plates at a seeding density of ~5000 cells/plate. Following overnight incubation in 5% CO_2_ incubator, 1 ml DMEM supplemented with 10% FBS was added to maintain hydration. After 2 weeks of growth, colonies were stained with 0.005% crystal violet/methanol for 1 h and subsequently photographed by light microscopy. The number of colonies in ~15–20 randomly-selected fields was counted under 10× magnification. Each experiment was repeated 3 times [biological repeats]; 3 technical repeats were included during each biological repeat.

### RNA isolation and qPCR—

These assays were carried out as described previously (Aznar et al, 2015). Total RNA was isolated using an RNeasy kit (QIAGEN) as per the manufacturers’ protocol. First-strand cDNA was synthesized using Superscript II reverse transcriptase (Invitrogen), followed by ribonuclease H treatment (Invitrogen) prior to performing quantitative real-time PCR. Reactions omitting reverse transcriptase were performed in each experiment as negative controls. Reactions were then run on a real-time PCR system (ABI StepOnePlus; Applied Biosystems). Gene expression was detected with SYBR green (Invitrogen), and relative gene expression was determined by normalizing to GAPDH using the ΔC_T_ method. Primer sequences are available upon request.

### Stratification of colon cancer patients in distinct gene-expression subgroups and comparative analysis of their survival outcomes—

The association between the levels of Daple (CCDC88C) and EGFR mRNA expression and patient survival was tested in cohort of 466 patients where each tumor had been annotated with the disease-free survival (DFS) information of the corresponding patient. This cohort included gene expression data from four publicly available NCBI-GEO data-series (GSE14333, GSE17538, GSE31595, GSE37892)(Jorissen et al, 2009; Laibe et al, 2012; Smith et al, 2010; Thorsteinsson et al, 2012), and contained information on 466 unique primary colon carcinoma samples, collected from patients at various clinical stages (AJCC Stage I-IV/Duke’s Stage A-D) by five independent institutions: 1) the H. Lee Moffit Cancer Center in Tampa, Florida, USA; 2) the Vanderbilt Medical Center in Nashville, Tennessee, USA; 3) the Royal Melbourne Hospital in Melbourne, Australia; 4) the Institut PaoliCalmette in Marseille, France; 5) the Roskilde Hospital in Copenhagen, Denmark. All 466 samples contained in this subset were cross-checked to exclude the presence of redundancies/duplicates. A complete list of all GSMIDs of the experiments contained within the NCBI-GEO discovery dataset has been published previously (Dalerba et al, 2016). To investigate the relationship between the mRNA expression levels of selected genes (i.e. CCDCDDC, Wnt5a, EGFR and FZD7) and the clinical outcomes of the 466 colon cancer patients represented within the NCBI-GEO discovery dataset, we applied the Hegemon, “hierarchical exploration of gene expression microarrays on-line” tool (Dalerba et al, 2011). The Hegemon software is an upgrade of the BooleanNet software (Sahoo et al, 2008), where individual gene-expression arrays, after having been plotted on a two-axis chart based on the expression levels of any two given genes, can be stratified using the StepMiner algorithm and automatically compared for survival outcomes using Kaplan-Meier curves and log-rank tests. Since all 466 samples contained in the dataset had been analyzed using the Affymetrix HG-U133 Plus 2.0 platform (GPL570), the threshold gene-expression levels for Daple/CCDC88C and EGFR were calculated using the StepMiner algorithm based on the expression distribution of the 25,955 experiments performed on the Affymetrix HG-U133 Plus 2.0 platform. We stratified the patient population of the NCBI-GEO discovery dataset in different gene-expression subgroups, based on either the mRNA expression levels of Daple/CCDC88C alone (i.e. CCDC88C neg vs. pos), EGFR alone (i.e. EGFR neg vs. pos), or a combination of both (i.e. CCDC88Cneg/EGFRpos vs. CCDC88Cpos/EGFRpos vs. CCDC88Cpos/EGFRneg vs CCDC88Cpos/EGFRpos). Once grouped based on their gene-expression levels, patient subsets were compared for survival outcomes using both Kaplan-Meier survival curves and multivariate analysis based on the Cox proportional hazards method.

### Statistical analysis—

Each experiment presented in the figures is representative of at least three independent experiments. Displayed images and immunoblots are representative of the biological repeats. Statistical significance between the differences of means was calculated by an unpaired student’s t-test or one-way ANOVA [whenever more than two groups were compared]. A two-tailed *p* value of <0.05 at 95% confidence interval is considered statistically significant. All graphical data presented were prepared using GraphPad or Matlab.

## ACKNOWLEDGMENTS

We thank Gordon N. Gill and Marilyn G. Farquhar (UCSD) and Deepali Bhandari (CSULB) for their critical input during the preparation of the manuscript. This work was supported by NIH grants CA100768, CA160911 and DK099226 (to P.G). P.G. was also supported by the American Cancer Society (ACS-IRG 70-002) and by the UC San Diego Moores Cancer Center. I.K. was supported by NIH grants GM071872, AI118985, and GM117424. I.L-S was supported by a fellowship from the American Heart Association (AHA #14POST20050025). F.H was supported by the State Scholarship Fund of China Scholarship Council (No.201208510048) and fund from West China Hospital, Sichuan University, PR China, during her tenure as a visiting professor to UCSD.

## COMPETING FINANCIAL INTERESTS

The authors declare no competing financial interests.

## AUTHOR CONTRIBUTIONS

N.A and P.G designed, performed and analyzed most of the experiments in this work. Y.D. cloned and generated all Daple mutants used in this work. N.S, F.H and K.S assisted with protein expression, purification, and in carrying out protein-protein interaction assays; I.L-S carried out in vitro kinase assays; M.G performed the mass spectrometry analyses. D.S and P.G analyzed the patient datasets using the Hegemon software; I.K generated the homology model for Daple-bound Dvl and provided structure-based guidance to study Daple mutants. N.A and P.G conceived the project and wrote the manuscript. P.G supervised and funded the project.

